# Cryo-EM Structures of Amyloid-β 42 Filaments from Human Brain

**DOI:** 10.1101/2021.10.19.464936

**Authors:** Yang Yang, Diana Arseni, Wenjuan Zhang, Melissa Huang, Sofia Lövestam, Manuel Schweighauser, Abhay Kotecha, Alexey G. Murzin, Sew Y. Peak-Chew, Jennifer Macdonald, Isabelle Lavenir, Holly J. Garringer, Ellen Gelpi, Kathy L. Newell, Gabor G. Kovacs, Ruben Vidal, Bernardino Ghetti, Benjamin Falcon, Sjors H.W. Scheres, Michel Goedert

**Author notes:** Equal contributions (Yang Yang, Diana Arseni, Wenjuan Zhang). MRC Prion Unit and Institute of Prion Diseases, University College, London, UK.

## Abstract

Filament assembly of amyloid-β peptides ending at residue 42 (Aβ42) is a central event in Alzheimer’s disease. We report the cryo-EM structures of Aβ42 filaments from brain. Two structurally related S-shaped protofilament folds give rise to two types of filaments. Type I filaments were found mostly in the brains of individuals with sporadic Alzheimer’s disease and Type II filaments in individuals with familial Alzheimer’s disease and other conditions. The structures of Aβ42 filaments from brain differ from those of filaments assembled *in vitro*. By contrast, in *App*^NL-F^ knock-in mice, Aβ42 deposits were made of Type II filaments. Knowledge of Aβ42 filament structures from human brain may lead to the development of inhibitors of assembly and improved imaging agents.

## Main Text

Alzheimer’s disease is defined by the simultaneous presence of two different filamentous amyloid inclusions in brain: abundant extracellular plaques of Aβ and intraneuronal neurofibrillary tangles of tau (*1*). Genetic evidence has indicated that Aβ is key to the pathogenesis of Alzheimer’s disease (*2*,*3*). Multiplications of the *APP* gene encoding the Aβ precursor protein, as well as mutations in *APP* and in *PSEN1* and *PSEN2*, the presenilin genes, cause familial Alzheimer’s disease. Presenilins form part of the γ-secretase complex that is required for the production of Aβ from *APP*. Although variability in γ-secretase cleavage results in Aβ peptides that vary in size, those of 40 (Aβ40) and 42 (Aβ42) amino acids are the most abundant. Mutations associated with familial Alzheimer’s disease increase the ratios of Aβ42 to Aβ40 (*4*,*5*), the concentration of Aβ42 (*6*) or the assembly of Aβ42 into filaments (*7*).

Three major types of Aβ inclusions are typical of the brain in Alzheimer’s disease (*8*–*11*): diffuse and focal deposits in the parenchyma, as well as vascular deposits. Diffuse deposits, which contain loosely packed Aβ filaments, are found in several brain regions, including entorhinal cortex, pre-subiculum, striatum, brainstem, cerebellum and subpial area. Focal deposits, in the form of dense core plaques, contain a spherical core of tightly packed Aβ filaments surrounded by more loosely packed filaments. Dense core plaques are found mostly in hippocampus and cerebral cortex. In advanced cases of Alzheimer’s disease, diffuse and focal Aβ deposits are widespread. In around 80% of cases of Alzheimer’s disease, Aβ deposits are also found in the walls of blood vessels (cerebral amyloid angiopathy). Electron cryomicroscopy (cryo-EM) provided the structures of Aβ40 aggregates from the meninges of Alzheimer’s disease brain (*12*). Meningeal deposits have a high Aβ40 and a low Aβ42 content and are morphologically distinct from parenchymal plaques.

Diffuse plaques and the loosely packed material of dense core plaques consist mainly of filamentous Aβ42, whereas plaque cores and blood vessel deposits are made of both Aβ40 and Aβ42. Aβ42 aggregates faster than Aβ40 and is the major species in plaques, despite the proteolytic processing of *APP* generating more soluble Aβ40 (*4*,*8*,*13*).

Aβ deposition appears to follow spatiotemporal spreading, suggesting that pathology may propagate through seeded aggregation, similar to prions (*14*–*16*). A prion-like mechanism may also explain the formation of Aβ deposits in cerebral blood vessels in some adults who received intramuscular injections of contaminated human growth hormone preparations as children and in individuals who were given dura mater grafts or underwent neurosurgery (*17*–*19*), even though they did not have the symptoms of Alzheimer’s disease. Besides Alzheimer’s disease, Aβ42 deposits can also be present as copathology in a number of other conditions, especially as a function of age (*10*). Despite their importance for disease pathogenesis, the structures of Aβ42 filaments from brain are not known.

Here we used cryo-EM to determine the structures of Aβ42 filaments extracted from the brains of ten individuals (Figs. 1, S1, Table S1). Five had Alzheimer’s disease, with three sporadic and two familial (mutation in *APP* encoding V717F and mutation in *PSEN1* encoding F105L) cases. Five individuals had other conditions, with a case of age-related tau astrogliopathy (ARTAG), a case of Parkinson’s disease dementia (PDD), a case of dementia with Lewy bodies (DLB), a case familial frontotemporal dementia (FTD) caused by a *GRN* mutation and a case of pathological aging (PA).

**Fig. 1.**
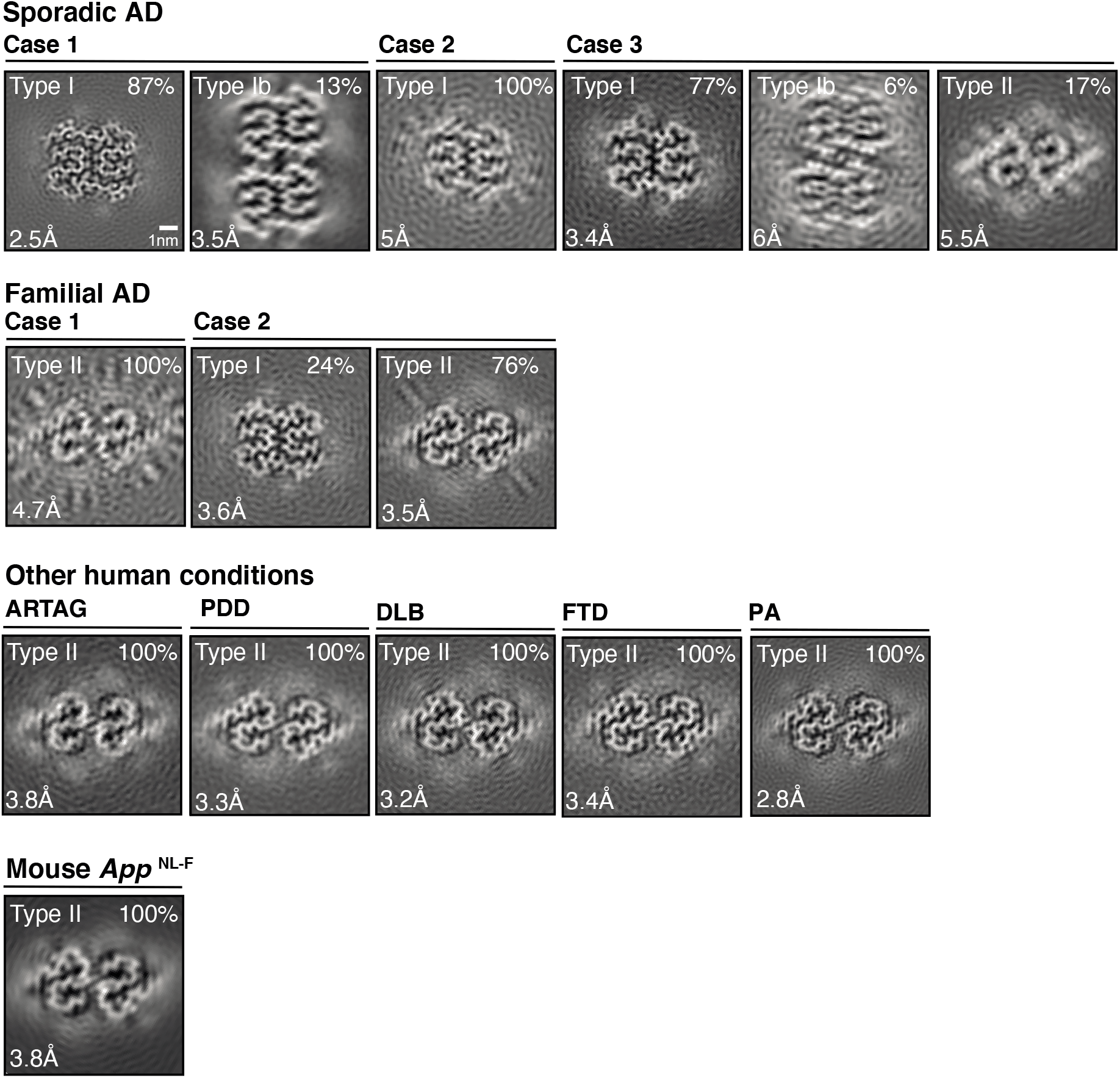
Cryo-EM maps of Type I, Type lb and Type II Aβ42 filaments from brain. Cryo-EM maps of Aβ42 filaments from five cases of Alzheimer’s disease [three sporadic (AD cases 1-3) and two familial (case 1, mutation in *APP*encoding V717F; case 2, mutation in *PSEN1* encoding F105L)]; other human diseases [a case of aging-related tau astrogliopathy (ARTAG), a case of Parkinson’s disease dementia (PDD), a case of dementia with Lewy bodies (DLB), a case of frontotemporal dementia (FTD) caused by a *GRN* mutation and a case of pathological aging (PA)]; and homozygous mice of the *App*^NL-F^ knock-in line. For each map, a sum of the reconstructed densities for several XY-slices, approximating one β-rung, is shown. Filament types (Type I, Type lb or Type II) are indicated on the top left; the percentages of a given filament type among Aβ filaments in the dataset are shown on the top right. The same scales apply to all panels.

### Type I Aβ42 Filaments from Human Brain

For individuals with sporadic Alzheimer’s disease, we observed a majority of twisted Aβ filaments, which we named Type I filaments (Figs. 1, 2A, B, D). They are made of two identical S-shaped (a double curve resembling the letter S or its reverse) protofilaments embracing each other with extended arms. The 2.5 Å resolution map of Type I filaments from sporadic Alzheimer’s disease case 1 was used to build the atomic model. The ordered core of each protofilament spans G9 to A42, with the S-shaped domain extending from residues 19 to 42 and the N-terminal arm comprising residues 9 to 18. The secondary structure of protofilaments comprises six short β-strands and intervening turns. The S-shaped domain folds around two hydrophobic clusters: the N-terminal part around the side chains of F19, F20, V24 and I31, and the C-terminal part around the side chains of A30, I32, M35, V40 and A42 (Fig. 2B, D).

**Fig. 2.**
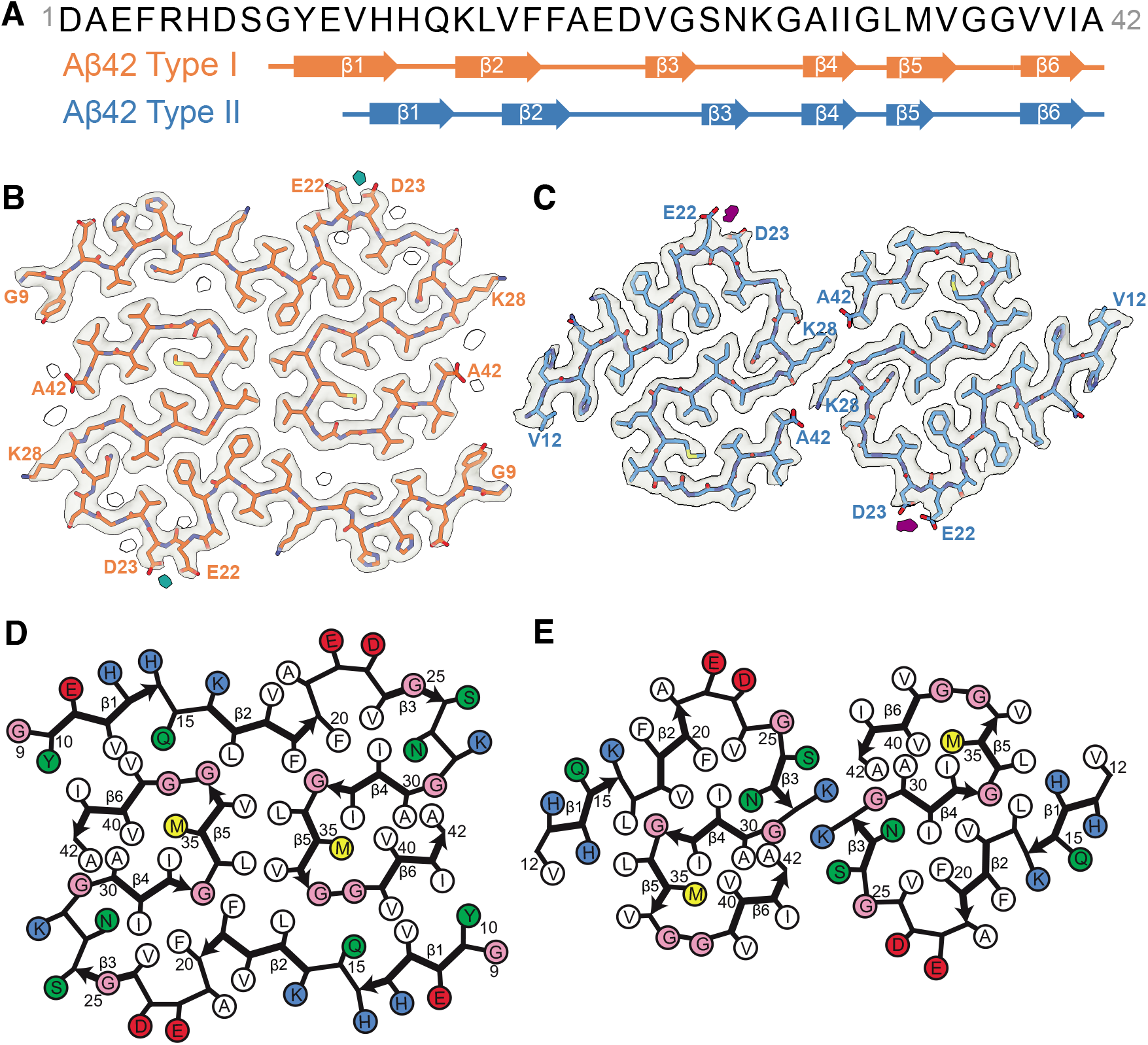
Structures of Type I and Type II Aβ42 filaments from brain. (A) Amino acid sequence of Aβ1-42. Type I filaments (in orange) extend from G9-A42; Type II filaments (in blue) from V12-A42. Thick connecting lines with arrowheads indicate β-strands (β1-β6). (B,C) Cryo-EM density map (in transparent grey) and atomic models for Type I (B) and Type II (C) Aβ42 filaments. Each filament type is made of two identical protofilaments shown in orange (Type I) or blue (Type II). Associated solvent molecules are shown in white and putative metals in teal (B) or purple (C). (D,E) Schematics of Type I (D) and Type II (E) Aβ42 folds. Negatively charged residues are shown in red, positively charged residues in blue, polar residues in green, apolar residues in white, sulfur-containing residues in yellow and glycines in pink. Thick connecting lines with arrowheads indicate β-strands.

The two protofilaments pack against each other with pseudo-21 symmetry (Fig. S2A). They form a predominantly hydrophobic interface involving the side chains of L34, V36, V39 and I41 on the outer surface of the S-shaped domain, and the side chains of Y10, V12, Q15 and L17 in the N-terminal arm. In sporadic Alzheimer’s disease cases 1 and 3, we also observed a minority of Type Ib filaments, in which two Type I filaments run side by side (Fig. 1).

Several additional densities, attributed to ordered solvent molecules, are resolved in the 2.5 Å resolution cryo-EM map (Figs. 2B, 3A). One of these, located adjacent to the negatively charged carboxyl groups of E22 and D23 on the filament surface, most likely corresponds to a bound metal ion (Fig. 3B, C). The conformations of both acidic residues are highly restrained: the side chain of E22 comes into close contact with the main chain of successive Aβ peptides along the helical axis, whereas the side chain of D23 is fastened to the main chain through hydrogen bonding. The binding of metal ions would alleviate the electrostatic repulsion between their negatively charged carboxyl groups, thus helping to stabilise the filament fold. These ions may, therefore, act as cofactors for filament formation (*20*). By contrast, there are no additional densities associated with an ordered grid of imidazole groups formed on the surface of Type I filaments by two adjacent histidine residues, H13 and H14. Their side chains are held together by a hydrogen bond, with H13 making a second hydrogen bond with the side chain of E11 in the next Aβ42 molecule.

**Fig. 3.**
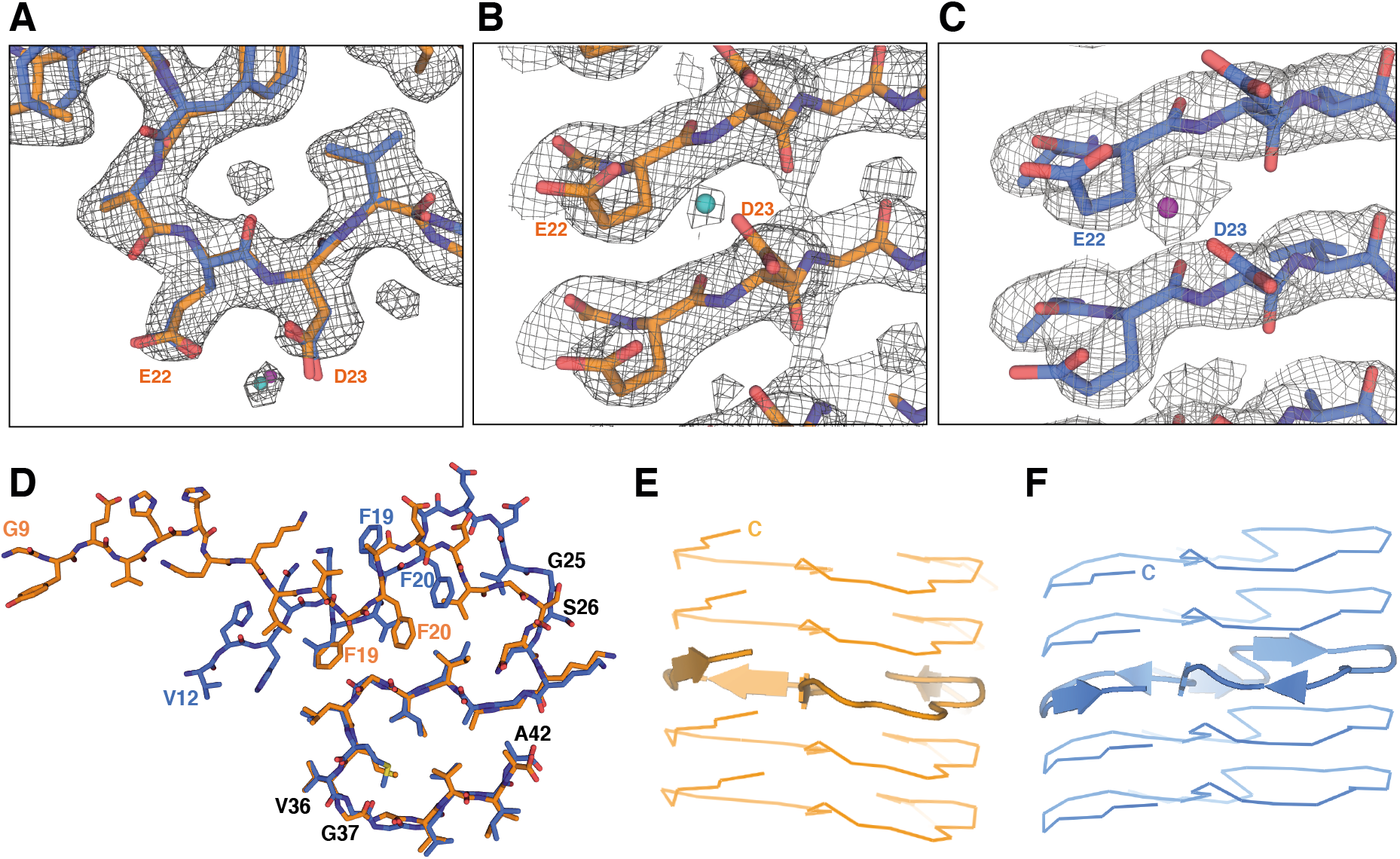
Protofilament folds and putative metal ion-binding sites of Type I and Type II Aβ42 filaments. (A) Superposition of the structures of F20-V24 arcs overlaid on the corresponding section of the 2.5 Å map of Type I filaments. Putative metal ions are shown as teal and purple spheres. (B,C) Side views of putative metal ion-binding sites in Type I (teal) and Type II (purple) protofilaments, superimposed on the corresponding density maps. (D) Superposition of Type I (orange) and Type II (blue) protofilaments, based on the central layer of their S-shaped domains. (E,F) Side views of Type I and Type II protofilaments along the central β4 strand. The centre layer monomers in five successive rungs are shown in cartoon, with β-strands shown as arrows.

### Type II Aβ42 Filaments from Human Brain

For individuals with familial Alzheimer’s disease and other conditions, we observed a major, twisted filament type, distinct from Type I, which we named Type II (Figs. 1, 2A, C, E). In case 3 of sporadic Alzheimer’s disease, 17% of filaments were Type II, whereas in case 2 of familial Alzheimer’s disease, 24% of filaments were Type I. The atomic model of Type II filaments, built using the 2.8 Å resolution map obtained for the case of PA, revealed that the ordered core extends from V12 to A42. Residues F20-A42 adopt a similar S-shaped fold to that of Type I filaments, with the same side chain orientations. Differences between folds are mostly limited to the orientations of a few peptide groups affecting secondary structure assignments. Peptides G25-S26 and V36-G37 are flipped by approximately 180° in the Type II fold. The flipped G25-S26 peptide results in a slight expansion of the N-terminal hydrophobic cluster by accommodating the side chains of L17 and V18 instead of F19, which faces outwards in Type II filaments (Fig. 3D). The reorientation of the second peptide leads to a shift of the C-terminal segment of the Type II fold along the helical axis by approximately one Aβ peptide, compared to its position in the Type I fold (Fig. 3E, F).

When compared to the Type I protofilament interface, that of Type II protofilaments is smaller and is formed by the opposite side of the S-shaped fold. Type II protofilaments pack against each other with C2 symmetry (Fig. S2B). The protofilament interface is primarily stabilised by electrostatic interactions between the amino group of K28 from one protofilament and the C-terminal carboxyl group of A42 from the other, and vice versa (Fig. 2C). Unlike Type I filaments, hydrophobic residues on the outer surfaces of the S-shaped domains remain exposed, forming non-polar patches on the surface of Type II filaments. There are fewer additional densities for ordered solvent molecules visible in the 2.8 Å map of Type II filaments than in the 2.5 Å map of Type I filaments, but the density for the putative metal ion bound to E22 and D23 is prominent in the equivalent location (Figs. 2B,C, 3E,F).

### Comparison with Known Structures

Type I and Type II filaments have a left-handed twist and are structurally different from Aβ40 aggregates from the meninges of individuals with Alzheimer’s disease, which comprise two identical protofilaments with an unrelated C-shaped fold and a right-handed twist (*12*). They also differ from the cryo-EM structures of left-handed Aβ40 filaments, which were derived from the cerebral cortex of an individual with Alzheimer’s disease by seeded filament growth (*21*), but share with them a common substructure (Fig. 4A). In the seeded Aβ40 filaments, which comprised two extended protofilaments, residues G25-G37 adopted virtually the same conformation as in the middle of the S-shaped fold of Type I and Type II filaments. Structures of Aβ42 filaments assembled *in vitro*, obtained by cryo-EM (*22*) and by solid-state NMR (*23*–*25*), each have a single or two identical protofilaments with an S-shaped domain like that of Type I and Type II filaments (Fig. 4B). In two NMR structures, the inter-protofilament packing also resembled that of Type I filaments. However, when examined at the residue level, none of the Aβ42 filaments assembled *in vitro* displayed the same side chain orientations and contacts or the same inter-protofilament packing as in Type I and Type II filaments. The structures of *in vitro* assembled filaments of Aβ40 with the Osaka mutation (deletion of codon 693 in *APP*, corresponding to E22 in Aβ), based on a large number of unambiguous intra- and intermolecular solid-state NMR distance restraints, are most similar to those of Type I Aβ42 filaments (Fig. 4C) (*26*).

**Fig. 4.**
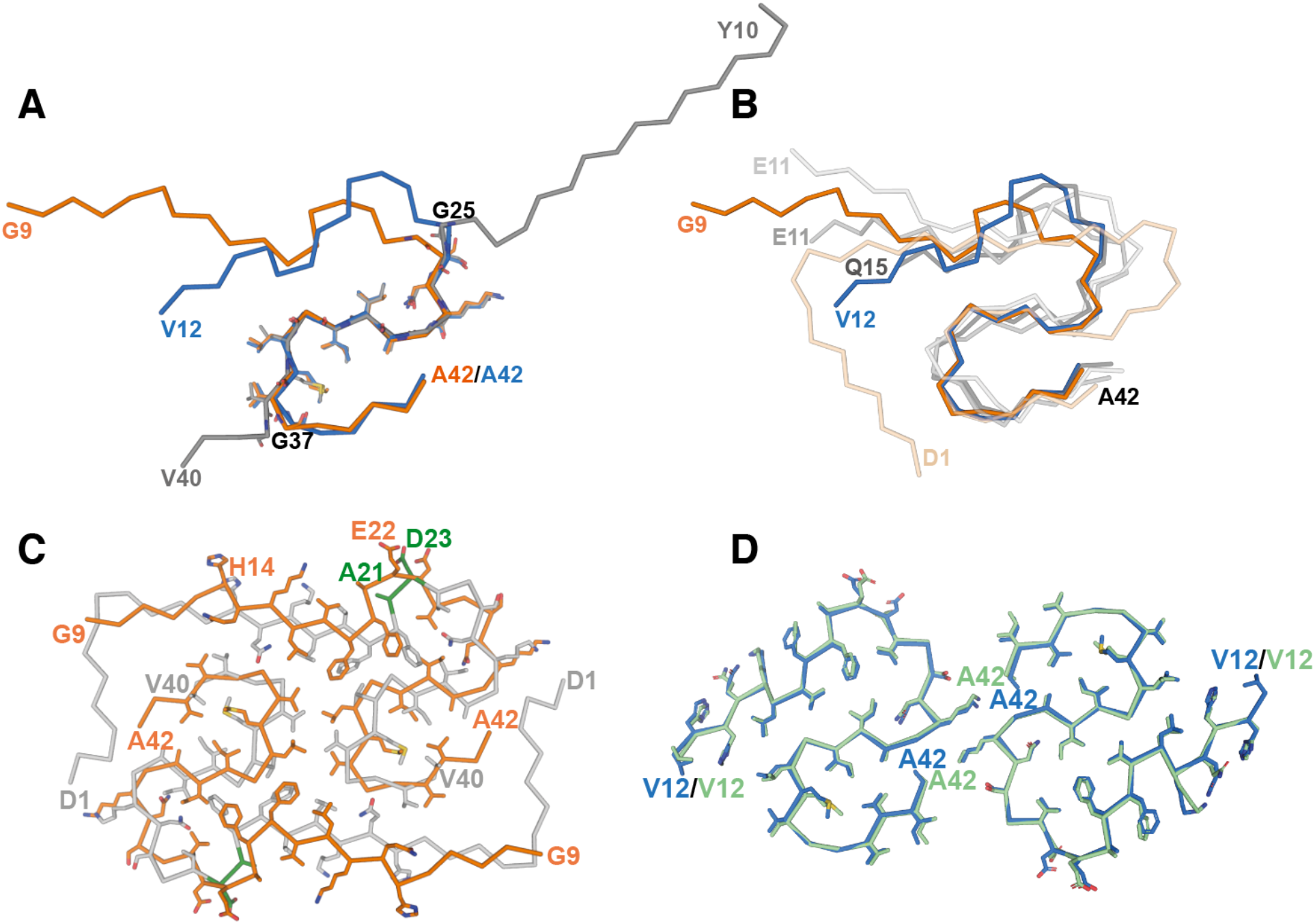
Comparison of protofilaments and filaments of brain Aβ42 with those of seeded recombinant Aβ40, recombinant Aβ42 and recombinant Aβ40ΔE22. (A) Comparison of cryo-EM structures of human brain Type I and Type II Aβ42 protofilaments with cryo-EM structure of seeded recombinant Aβ40 protofilaments. Type I is in orange; Type II is in blue; seeded Aβ40 (PDB 6W0O) is in grey. (B) Comparison of cryo-EM structures of human brain Type I and Type II Aβ42 protofilaments with cryo-EM and NMR structures of recombinant Aβ42 protofilaments. PDBs and colour codes for recombinant Aβ42 filaments: 5OQV, wheat; 2NAO, dark grey; 2MXU, grey; 5KK3, light grey. (C) Comparison of cryo-EM structures of human brain Type I Aβ42 filaments and NMR structures of recombinant Aβ40ΔE22 filaments. Human brain Type I is in orange; recombinant Aβ40ΔE22 is in grey, with residues around ΔE22 shown in green. (D) Comparison of cryo-EM structures of Aβ42 filaments from the brains of mouse knock-in line *AppF*^NL-F^ with human brain Type II Aβ42 filaments. Mouse brain filaments are in green; human brain Type II filaments are in blue.

Reconstructions of Type I and Type II filaments show strong densities for residues 41 and 42, indicating that the majority of molecules corresponds to Aβ42. In agreement, Western blotting (Fig. S3) and mass spectrometry (Fig. S4) of the extracted filaments showed that Aβ42 was the majority species in all cases, with variable amounts of Aβ40.

We performed immunohistochemistry on the contralateral sides of the brain regions used for cryo-EM, immunoblotting and mass spectrometry (Figs. S5, S6). Deposits of Aβ42 were also more numerous than those of Aβ40, with sporadic Alzheimer’s disease cases 1 and 3, familial Alzheimer’s disease case 1, as well as the cases of ARTAG, PDD, DLB and PA showing almost exclusively Aβ42 deposits. Most deposits of Aβ40 were present in sporadic and familial Alzheimer’s disease cases 2. By immunohistochemistry, Aβ40 deposits were also abundant in FTD. This difference with immunoblotting and mass spectrometry may reflect a hemispheric asymmetry in Aβ deposition. Plaque cores were most numerous in sporadic Alzheimer’s disease cases 1-3 and blood vessel deposits of Aβ40 were found in sporadic and familial Alzheimer’s disease cases 2. In all cases, diffuse deposits of Aβ were more numerous than focal and blood vessel deposits.

It is possible that low levels of Aβ40, or shorter peptides, may be incorporated in Type I and Type II filaments. The inter-protofilament salt bridge between K28 of one protofilament and the C-terminal carboxyl of A42 of the other, in Type II, but not Type I, filaments, suggests that it is more likely that hybrid Aβ42/Aβ40 filaments will be of Type I. This is supported by the structural similarities of Type I Aβ42 filaments with filaments of Aβ40 with the Osaka mutation (Fig. 4C) (*26*). We did not find evidence for filaments composed predominantly of Aβ40. However, we cannot exclude that such filaments were present in low amounts, or were not extracted as dispersed filaments suitable for cryo-EM reconstruction.

Depending on the filament type, 8 or 11 residues are disordered at the amino-terminus. The β-site *APP* cleaving enzyme 1 (BACE1) generates the amino-terminus of Aβ (*27*). BACE1 mainly cleaves at residue 1 of Aβ, but some cleavage at residues 11 or 12 also occurs. Structures of Type I and Type II filaments from brain are compatible with the incorporation of shorter peptides. However, by Western blotting and mass spectrometry (Figs. S3, S4), the bulk of Aβ42 in the extracted filaments was full-length. It follows that the amino-terminal residues that are not present in Type I or Type II filament cores form the fuzzy coat of Aβ42 filaments. This is supported by the decoration of Type I and Type II filaments using antibodies specific for the amino-terminal region of Aβ (Fig. S7). Tau filaments were unlabelled. The fuzzy coat of Aβ42 filaments thus comprises around 20% of the molecule, with the core making up 80%. By contrast, the fuzzy coat of tau filaments from Alzheimer’s disease amounts to over 80% (*28*).

*In vitro* aggregation is essential for studying the molecular mechanisms that underlie amyloid formation. However, available methods for the assembly of recombinant tau and α-synuclein yield filament structures that are different from those of filaments extracted from human brain (*29*–*32*). The same appears to be true of Aβ42 filaments, which only partially reproduce the structures from human brain (*29*–*32*).

### Type II Aβ42 Filaments from *App*^NL-F^ Mouse Brain

Animal models provide another tool for studying the molecular mechanisms of Alzheimer’s disease. *App*^NL-F^ knock-in mice express humanized Aβ and harbour the Swedish double mutation (KM670/671NL), as well as the Beyreuther/Iberian mutation (I716F) in *App* (*33*). They develop abundant deposits of wild-type human Aβ42, neuroinflammation and memory impairment, without requiring the overexpression of *APP*. To further study the relevance of this mouse model for human disease, we determined the cryo-EM structures of Aβ42 filaments from the brains of 18-month-old homozygous *App*^NL-F^ mice (Figs. 1, 4D). They were identical to Type II filaments from human brain, making this mouse line the first known experimental system with filament structures like those from human brain. It is possible that cofactors required for the formation of Type II filaments are present in the brains of *App*^NL-F^ knock-in mice, but missing in *in vitro* experiments.

## Discussion

Type I and Type II Aβ42 filaments from brain are each made of two identical protofilaments, but the protofilaments of Type I filaments differ from those of Type II. This is unlike tau assembly in human brain, where a single protofilament gives rise to two or more types of filaments (*34*) and α-synuclein in multiple system atrophy, where four protofilaments give rise to two different filaments (*31*). Here, Type I filaments were limited to cases of sporadic Alzheimer’s disease that had also the largest number of plaque cores. A majority of Type II filaments was present in cases with abundant diffuse deposits of Aβ and a smaller number of focal plaques with cores. This included cases of familial Alzheimer’s disease, as well as cases of ARTAG, PDD, DLB, FTD and PA. Cases of Alzheimer’s disease with a majority of Type I filaments were older at death than other Alzheimer and non-Alzheimer cases with a majority of Type II filaments in neocortex. There was no correlation between Aβ42 filament type and *APOE* genotype. The relevance of these differences between Type I and Type II filaments is not known. Because positron emission tomography compound PiB (Pittsburgh compound B) visualizes Aβ deposits in both sporadic and familial cases of Alzheimer’s disease, it probably labels both filament types (*35*).

Like V717F, the mutation in *APP* encoding I716F, increases the ratio of Aβ42 to Aβ40 (*4*,*36*,*37*). This may explain the presence of Type II Aβ42 filaments in *App*^NL-F^ mice and in human cases with F717 *APP*. Line *App*^NL-F^ may therefore be a model for some cases of familial Alzheimer’s disease, but not necessarily of sporadic disease.

Differential labelling by luminescent conjugated oligothiophene amyloid ligands suggested substantial heterogeneity in the molecular architecture of Aβ deposits from the brains of patients with Alzheimer’s disease (*38*). Our findings indicate that this heterogeneity is not the result of differences in the structures of Aβ42 filaments. As suggested for Aβ40 (*19*,*39*), a single Aβ42 filament type predominated in a given Alzheimer’s disease brain. Together with a second filament type, it accounted for the Aβ42 filaments from different cases of Alzheimer’s disease, ARTAG, PDD, DLB, FTD and PA. Knowledge of the structures of Aβ42 filaments from brain may lead to the development of better *in vitro* and animal models for these diseases, inhibitors of Aβ42 assembly and imaging agents with increased specificity and sensitivity.

## Acknowledgments

We thank the patients’ families for donating brain tissues; U. Kuederli, M. Jacobsen, F. Epperson and R.M. Richardson for human brain collection and technical support; T. Saido for providing *App*^NL-F^ mice; T. Darling and J. Grimmett for help with high-performance computing; G. Singla Lezcano for help with Falcon 4i; Y. Shi, J. Collinge and C. Haass for helpful discussions. This study was supported by the Electron Microscopy Facility of the MRC Laboratory of Molecular Biology.

## Author contributions

E.G.,K.L.N., G.G.K. and B.G. identified patients and performed neuropathology; H.J.G. and R.V. performed genetic analysis; J.M., I.L. and M.H. organized breeding and characterized mouse tissues; Y.Y., D.A., W.Z. and S.Y.P.-C. prepared Aβ filaments and performed Western blotting and mass spectrometry; Y.Y., D.A., W.Z., M.H. and M.S. performed cryo-EM data acquisition; Y.Y., D.A., W.Z., M.H., S.L., A.K. A.G.M., B.F. and S.H.W.S. performed cryo-EM structure determination; B.F., S.H.W.S. and M.G. supervised the project; all authors contributed to writing the manuscript.

## Data and materials availability

There are no restrictions on data availability and data are deposited…

## Supplementary Materials

Materials and Methods

Tables S1-S3 Figures S1-S7

References (*40*-*62*)

## Material and Methods

### Clinical history and neuropathology

We determined the cryo-EM structures of Aβ42 filaments from the brains of ten individuals (Table S1). Five had Alzheimer’s disease [three sporadic and two familial (mutation in *APP* encoding V717F and mutation in *PSEN1* encoding F105L) cases]. Five individuals had other conditions: a case of aging-related tau astrogliopathy (ARTAG); a case of Parkinson’s disease dementia (PDD); a case of dementia with Lewy bodies (DLB); a case of frontotemporal dementia (FTD) caused by a *GRN* mutation; and a case of pathological aging (PA), defined as abundant Aβ deposits in neocortex with no or only limited tau pathology, in the absence of dementia (frontal cortex) (*40*). *APP*, *PSEN-1* and *PSEN-2* were sequenced; with the exception of familial Alzheimer’s disease cases 1 and 2, sequences were wild-type (Table S1). *APOE* genotypes are also listed in Table S1.

Case 1 of sporadic Alzheimer’s disease was a 79-year-old man who died with a neuropathologically confirmed diagnosis following an 8-year history of progressive dementia. The frontal lobes were mildly atrophic. Abundant neurofibrillary tangles and neuritic plaques were present in frontal cortex. Case 2 of sporadic Alzheimer’s disease was a 82-year-old woman who died with a neuropathologically confirmed diagnosis following a 9-year history of progressive dementia. The frontal lobes were moderately atrophic. Abundant neurofibrillary tangles and neuritic plaques were observed in the frontal cortex and moderate Aβ angiopathy was present in the leptomeninges. We described this case before: number 2 in (*28*). Case 3 of sporadic Alzheimer’s disease was an 80-year-old woman who died with a neuropathologically confirmed diagnosis after a 9-year history of progressive dementia. The parietal lobes were mildly atrophic. Abundant neuritic plaques and moderate to severe neurofibrillary tangles were present in parietal cortex. Mild Aβ angiopathy was observed in parenchymal blood vessels of the parietal cortex. α-Synuclein deposits were present in the amygdala, within nerve cell bodies and processes. Case 1 of familial Alzheimer’s disease was a 54-year-old woman who died with a neuropathologically confirmed diagnosis following a 9-year history of progressive dementia caused by a mutation in *APP* encoding V717F. Her sister, mother and maternal grandfather had suffered from dementia caused by the same mutation. They belonged to the family described in (*41*). The frontal lobes were severely atrophic and the lateral ventricles were enlarged, especially in frontal and occipital horns. Nerve cell loss and gliosis were severe throughout the neocortex. Abundant diffuse Aβ plaques were in evidence. Moderate to severe neuritic plaques and neurofibrillary tangles were present in the frontal cortex. Moderate Aβ angiopathy was observed. α-Synuclein deposits were present in nerve cell bodies and processes of the amygdala. We described this case before: number 16 in (*28*). Case 2 of familial Alzheimer’s disease was a 67-year-old woman who died with a neuropathologically confirmed diagnosis following an 11-year history of progressive dementia caused by a *PSEN1* mutation encoding F105L. Her father and two siblings suffered from dementia caused by the same mutation; a sibling is described in (*42*). The frontal lobes were severely atrophic, with many plaques and tangles, and severe nerve cell loss. Abundant diffuse Aβ deposits were in evidence, alongside smaller numbers of cored plaques and mild cerebral angiopathy. Cotton wool plaques were not a major feature.

ARTAG was seen in an 85-year-old woman with a 1-year history of cancer and depression (*43*). Upon neuropathological examination, she had prominent ARTAG (subpial, subependymal, grey matter, white matter and perivascular). In entorhinal cortex/hippocampus, abundant Aβ plaques (diffuse and cored) were present, alongside α-synuclein-positive Lewy pathology and TDP-43 inclusions. PDD was seen in a 64-year-old man with a 25-year history of Parkinson’s disease and a 10-year history of progressive cognitive decline. The frontal lobes were mildly atrophic. Abundant α-synuclein-positive Lewy bodies and Lewy neurites were present in the amygdala. Some Aβ plaques and tau tangles were also present. DLB was seen in a 73-year-old man with a 5-year history of progressive cognitive decline. Upon neuropathological examination, abundant α-synuclein-positive Lewy bodies and Lewy neurites were present in the temporal cortex. Frequent Aβ plaques were also in evidence. FTD was seen in a 66-year-old woman with a 5-year history of progressive cognitive decline, dysarthria and emotional lability, as well as an intronic (a) to (g) mutation two bases before the start of the coding region of exon 11 of *GRN*. This mutation is predicted to disrupt the normal splicing of exon 11. Three siblings and her mother died with dementia in their 60s and 70s. Frontotemporal lobar degeneration with abundant TDP-43-immunoreactive neuronal cytoplasmic inclusions and dystrophic neurites were in evidence. Moreover, diffuse Aβ plaques were found in multiple brain regions, including the frontal cortex. PA was seen in a 59-year-old non-demented man who died of cardiac arrest. He had moderate numbers of non-cored neuritic plaques and diffuse Aβ deposits, with a smaller number of cored deposits, in the absence of cerebral amyloid angiopathy. Tau tangles were not detected in frontal cortex.

### Mice

Homozygous *App*^NL-F^ knock-in mice (*33*) were maintained on a C57BL/6 background. They developed increasing numbers of neocortical Aβ42 deposits from 6 months of age onwards. We sacrificed 18-month-old mice by cervical dislocation and pooled the brains from three mice for the extraction of sarkosyl-insoluble Aβ filaments.

### Extraction of Aβ filaments

For cryo-EM, Western blotting and mass spectrometry, sarkosyl-insoluble material was extracted from frontal cortex (cases 1, 2, 4 and 5 of Alzheimer’s disease; DLB; FTD; pathological aging); parietal cortex (case 3 of Alzheimer’s disease); entorhinal cortex/hippocampus (ARTAG); amygdala (PDD), essentially as described (*31*). Briefly, tissues were homogenized in 20 vol (w/v) extraction buffer consisting of 10 mM Tris-HCl, pH 7.5, 0.8 M NaCl, 10% sucrose and 1 mM EGTA. Homogenates were brought to 2% sarkosyl and incubated for 60 min at 37° C. Following a 10 min centrifugation at 10,000 g, the supernatants were spun at 100,000 g for 60 min. The pellets were resuspended in 1 ml/g extraction buffer and centrifuged at 3,000 g for 5 min. The supernatants were diluted 3-fold in 50 mM Tris-HCl, pH 7.5, containing 0.15 M NaCl, 10% sucrose and 0.2% sarkosyl, and spun at 100,000 g for 30 min. Sarkosyl-insoluble pellets were resuspended in 100 μl/g of 20 mM Tris-HCl, pH 7.4, 50 mM NaCl and used for cryo-EM, Western blotting and mass spectrometry. By immuno-EM, numerous Aβ42 and tau filaments were in evidence. Only tau filaments were previously observed, when a standard sarkosyl extraction method was used (*28*,*29*). A major difference is that we here added sarkosyl following tissue homogenization.

### Immunoblotting and immunohistochemistry

For immunoblotting, sarkosyl-insoluble pellets were resuspended in fluorescence-compatible buffer (Invitrogen LC2570) containing 4% 2-mercaptoethanol, sonicated for 5 min at 50% amplitude (QSonica) and boiled for 8 min. They were resolved using 10-20% Tricine gels (Invitrogen) and Western blotting was done essentially as described (*44*). Primary antibodies were: Rabbit monoclonal antibody specific for Aβ40 (1:2,000, BioLegend 867802) and rabbit monoclonal antibody specific for Aβ42 (1:2,000, BioLegend 812101. Secondary antibodies were fluorescently labelled. Fluorescence was detected on a ChemiDoc MP system (Biorad). Immunohistochemistry was carried out as described (*44*). Brain sections were 8 μm thick and were counterstained with haematoxylin. Primary antibodies for Aβ40 (1:100) and Aβ42 (1:500) were the same as for immunoblotting.

### Mass spectrometry

Mass spectrometry was performed as described (*45*,*46*). Sarkosyl-insoluble pellets were resuspended in 100 μl HFIP (hexafluoroisopropanol, 1,1,1,3,3,3 hexafluoro-2-propanol). Following a 3 min sonication at 50% amplitude (QSonica), they were incubated at 37°C for 2 h and centrifuged at 100,000 g for 15 min, before being dried by vacuum centrifugation. The samples were then resuspended in 2% acetonitrile/2% formic acid and centrifuged at 13,000 rpm for 5 min. The supernatants were desalted using home-made C18 (3M Empore) stage tips containing 1.5 μl Poros R3 resin (Applied Biosystems). Bound peptides were eluted sequentially with 20 μl of 30%, 50% and 80% acetonitrile in 0.5% formic acid. Eluates were partially dried by vacuum centrifugation. Matrix-assisted laser desorption/ionization time of flight (MALDI-TOF) mass spectrometry measurements were performed using an Ultraflex III (Bruker Daltonics, Bremen) mass spectrometer in positive and reflection modes; 0.6 μl sample was spotted onto the MALDI-TOF target, followed by 0.6 μl α-cyano-4-hydroxycinnamic acid (CHCA) as matrix. Mass spectrometry data were searched using Mascot (Matric Science, v 2.4). For liquid chromatography with tandem mass spectrometry (LC-MS/MS), partially dried samples were diluted with 0.1% formic acid and separated using an ultimate 3,000 RSLC Nanosystem (Thermo Fisher), with an acetonitrile gradient at a flow rate of 300 nl/min. Eluted peptides were introduced into a Q exactive plus hybrid quadrupole-Orbitrap mass spectrophotometer (Thermo Fisher). The raw data files were searched using Mascot. Modifications were set as Glu to pyroGlu and oxidation of Met. Scaffold (version 4.8.4, Proteome Software, Inc) was used to validate MS/MS-based peptide identifications.

### Whole exome sequencing

Target enrichment made use of the SureSelectTX human all-exon library (V6, 58 mega base pairs; Agilent) and high-throughput sequencing was carried out using a HiSeq4,000 (2×75-base-pair paired-end configuration; Illumina). Bioinformatics analyses were performed as described (*47*).

### Immunogold negative-stain electron microscopy

Extracted Aβ filaments were deposited on glow-discharged 400 mesh formvar/carbon fibre-coated copper grids (EM Sciences CF400-Cu) for 40s, blocked for 10 min with PBS+0.1% gelatin and incubated with primary antibody (1:50) in blocking buffer, as described (*48*). Primary antibodies were: D54D2 (Cell Signalling Technology) and 1E11 (BioLegend), two monoclonal antibodies specific for the N-terminal region of Aβ. Following incubation with primary antibodies, the grids were rinsed with blocking buffer and incubated with 10 nm gold-conjugated anti-rabbit IgG (Sigma), diluted 1:20 in blocking buffer. They were then washed with water and stained with 2% uranyl acetate for 40s. Images were acquired as described (*20*).

### Electron cryo-microscopy

For cryo-EM, with the exception of *App*^NL-F^ mice, extracted Aβ42 filaments were centrifuged at 3,000 g for 2 min and treated with 0.4 mg/ml pronase for 30-60 min (*48*). Aliquots of 3 μl were applied to glow-discharged holey carbon grids (Quantifoil Au R1.2/1.3, 300 mesh) and plunge-frozen in liquid ethane using a Thermo Fisher Vitrobot Mark IV. For all cases, datasets were acquired on Thermo Fisher Titan Krios G2 or G3 microscopes, with Gatan K2 or K3 detectors in counting mode, using a Bio-quantum energy filter (Gatan) with a slid width of 20 e^−^V. Mouse data were acquired on a Thermo Fisher Titan Krios G2 microscope using a Falcon-4 detector and no energy filter. Further details are given in Fig. 1 and Table S2. For sporadic Alzheimer’s disease case 1, an additional dataset was recorded on a Thermo Fisher Titan Krios G4 microscope equipped with a cold field-emission gun, a Selectris X energy filter and a Falcon-4i detector. The energy filter was operated with a slit width of 10 e^−^V to remove inelastically scattered electrons. Images were recorded with a flux of 8.1 e^−^/pixel per second and a total dose of 40 e^−^/Å^2^. Data were acquired with aberration-free image shift (AFIS) using EPU software at a throughput of 573 images per hour.

### Helical reconstruction

All super-resolution frames were gain-corrected, binned by a factor of 2, aligned, dose-weighted and then summed into a single micrograph using RELION’s own implementation of MotionCor2 (*49*). Contrast transfer function (CTF) parameters were estimated using CTFFIND-4.1 (*50*). All subsequent image-processing steps were performed using helical reconstruction methods in RELION (*51*,*52*). Filaments were picked manually. Reference-free 2D classification was performed to identify homogeneous segments for further processing. Initial 3D reference models were reconstructed *de novo* from 2D class averages (*53*) using an estimated rise of 4.75 Å and helical twists according to the observed cross-over distances of the filaments in the micrographs for the datasets on Alzheimer’s disease cases 1 and 3, PDD and DLB. Refined models from these cases, low-pass filtered to 10 Å were used as initial models for the remaining cases. 3D classification was used to select the best particles from each dataset. In order to increase the resolution of the reconstructions, Bayesian polishing (*54*) and CTF refinement (*55*) were performed for all datasets. Final 3D reconstructions, after 3D auto-refinement, were sharpened using the standard post-processing procedures in RELION, and overall final resolutions were calculated from Fourier shell correlations at 0.143 between the two independently refined half-maps, using phase-randomisation to correct for convolution effects of a generous, soft-edged solvent mask (*56*). Further details of data acquisition and processing are given in Table S2. For sporadic Alzheimer’s disease case 1, the best map for Type I filaments at a resolution of 2.5 Å, was obtained from the additional dataset acquired on the Krios G4. The best map for Type Ib filaments, at 3.5 Å resolution, was obtained from the other dataset. The ratio of Type I and Type Ib filaments was the same in both datasets. The best maps are shown in Fig. 1.

### Model building and refinement

Atomic models were only built and refined in the best available maps, i.e. the Type I structure in the map for Alzheimer’s disease case 1 (collected on Krios G4) and the Type II and Type Ib filaments structure in the map for the case of pathological aging (collected on Krios G3). Initial model building started by fitting the S-shaped protofilaments of the Aβ42 Type I and Type II reconstructions. The handedness of the maps was deduced from two observations. First, in the map of the Type I filament from Alzheimer’s disease case 1 at 2.5 Å resolution, the chirality of individual amino acid residues was clearly discernable due to the densities for carbonyl oxygen atoms in the main chain, which were also visible for most residues in the map of the Type II filament from the case of pathological aging at 2.8 Å resolution. Secondly, residues 26-33 in Type I and Type II maps adopted the same conformation, including a chiral (left-handed) turn of the main chain, as in a 1.1 Å micro-electron diffraction structure of an Aβ20-34 peptide (*57*). The same conformation was also reported for a cryo-EM structure of Aβ1-40, with a left-handed twist to a resolution of 2.8 Å (*23*). Atomic models were built manually using COOT (*58*). Side chain clashes were detected using MOLPROBITY (*59*) and corrected by iterative cycles of real-space refinement in COOT and Fourier-space refinement in REFMAC (*60*) and/or real-space refinement in PHENIX (*61*). For each refined structure, separate model refinements were performed against a single half-map, and the resulting model was compared to the other half-map to confirm the absence of overfitting. Molecular graphics and analyses were performed in ChimeraX (*62*). Statistics for the final models are given in Table S3.

## Supplementary Tables

**Table S1.** Summary of human cases. Brain regions studied by cryo-EM are indicated, as are *APOE* genotypes. Exons encoding amyloid precursor protein (*APP*), presenilin 1 (*PSEN1*) and presenilin 2 (*PSEN2*) were sequenced.

| Conditions | Case number | Gender | Age at death (y) | Brain region | *APOE* | *APP* | *PSEN1* | *PSEN2* |
| --- | --- | --- | --- | --- | --- | --- | --- | --- |
| Sporadic Alzheimer's disease | 1 | M | 79 | Frontal cortex | $\epsilon 2/\epsilon 3$ | *WT* | *WT* | *WT* |
| | 2 | F | 82 | Frontal cortex | $\epsilon 4/\epsilon 4$ | *WT* | *WT* | *WT* |
| | 3 | F | 80 | Parietal cortex | $\epsilon 3/\epsilon 4$ | *WT* | *WT* | *WT* |
| Familial Alzheimer's disease | 1 | F | 54 | Frontal cortex | $\epsilon 3/\epsilon 3$ | *V717F* | *WT* | *WT* |
| | 2 | F | 67 | Frontal cortex | $\epsilon 2/\epsilon 3$ | *WT* | *F105L* | *WT* |
| Aging-related tau astrogliopathy | 1 | F | 85 | Limbic region | $\epsilon 3/\epsilon 3$ | *WT* | *WT* | *WT* |
| Parkinson's disease dementia | 1 | M | 64 | Amygdala | $\epsilon 3/\epsilon 4$ | *WT* | *WT* | *WT* |
| Dementia with Lewy bodies | 1 | M | 73 | Frontal cortex | $\epsilon 3/\epsilon 4$ | *WT* | *WT* | *WT* |
| Frontotemporal dementia | 1 | F | 66 | Frontal cortex | $\epsilon 3/\epsilon 3$ | *WT* | *WT* | *WT* |
| Pathological aging | 1 | M | 59 | Frontal cortex | $\epsilon 3/\epsilon 3$ | *WT* | *WT* | *WT* |

**Table S2.**
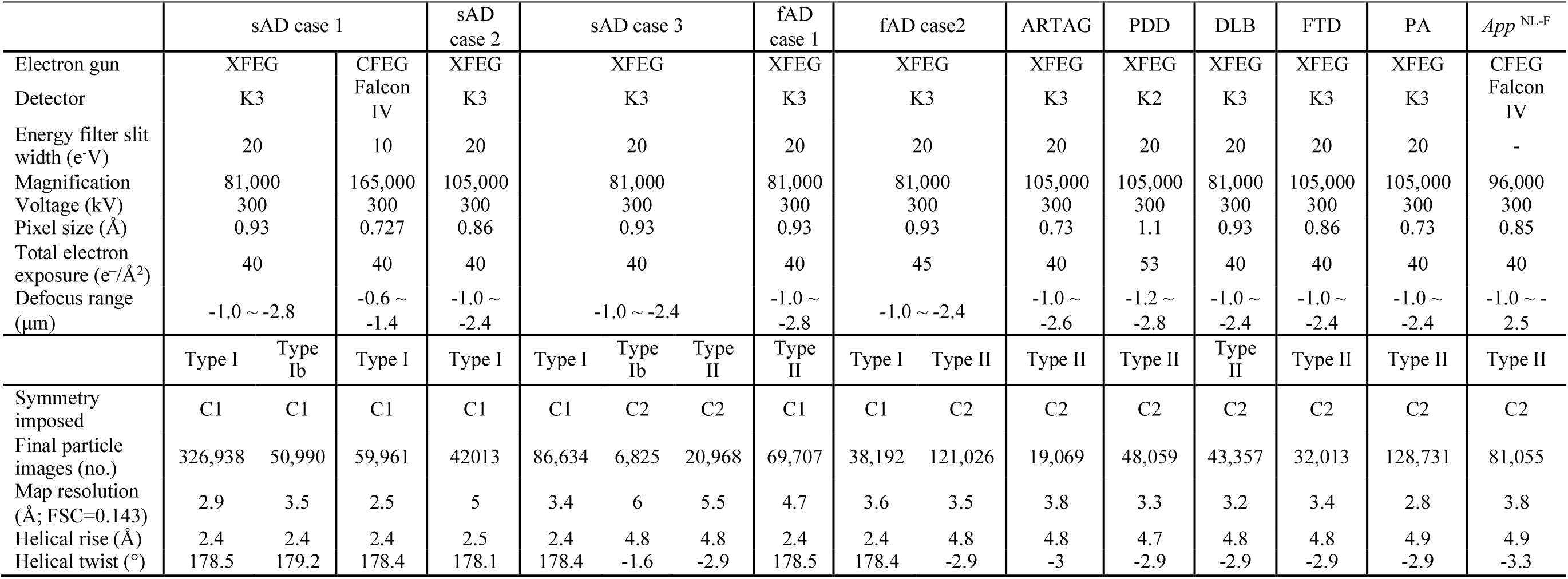
Statistics of data acquisition and processing.

**Table S3.** Refinement and model statistics of Type I and Type II Aβ42 filaments.

|  | sAD case 1 | PA |
| --- | --- | --- |
| **Model Composition** |  |  |
| Chains | 11 | 10 |
| Non-hydrogen atoms | 2,573 | 2,260 |
| Protein residues | 340 | 310 |
| Water | 60 | 0 |
| **Refinement** |  |  |
| Resolution (Å) | 2.5 | 2.8 |
| CC (mask) | 0.9 | 0.9 |
| **R.M.S deviations** |  |  |
| Bond lengths (Å) | 0.01 | 0.007 |
| Bond angles (°) | 0.71 | 0.79 |
| **Validation** |  |  |
| Molprobity score | 1.5 | 1.6 |
| Clashscore, all atoms | 5.4 | 3.9 |
| Rotamers outliers (%) | 0 | 0 |
| Cβ outliers (%) | 0 | 0 |
| **Ramachandran plot** |  |  |
| Favored (%) | 96.9 | 93.1 |
| Allowed (%) | 3.1 | 6.9 |
| Outliers (%) | 0 | 0 |
| PDB accession code | - | - |
| EMDB accession code | - | - |
| EMPIAR accession code | - | - |

## Supplementary Figures

**Fig. S1.**
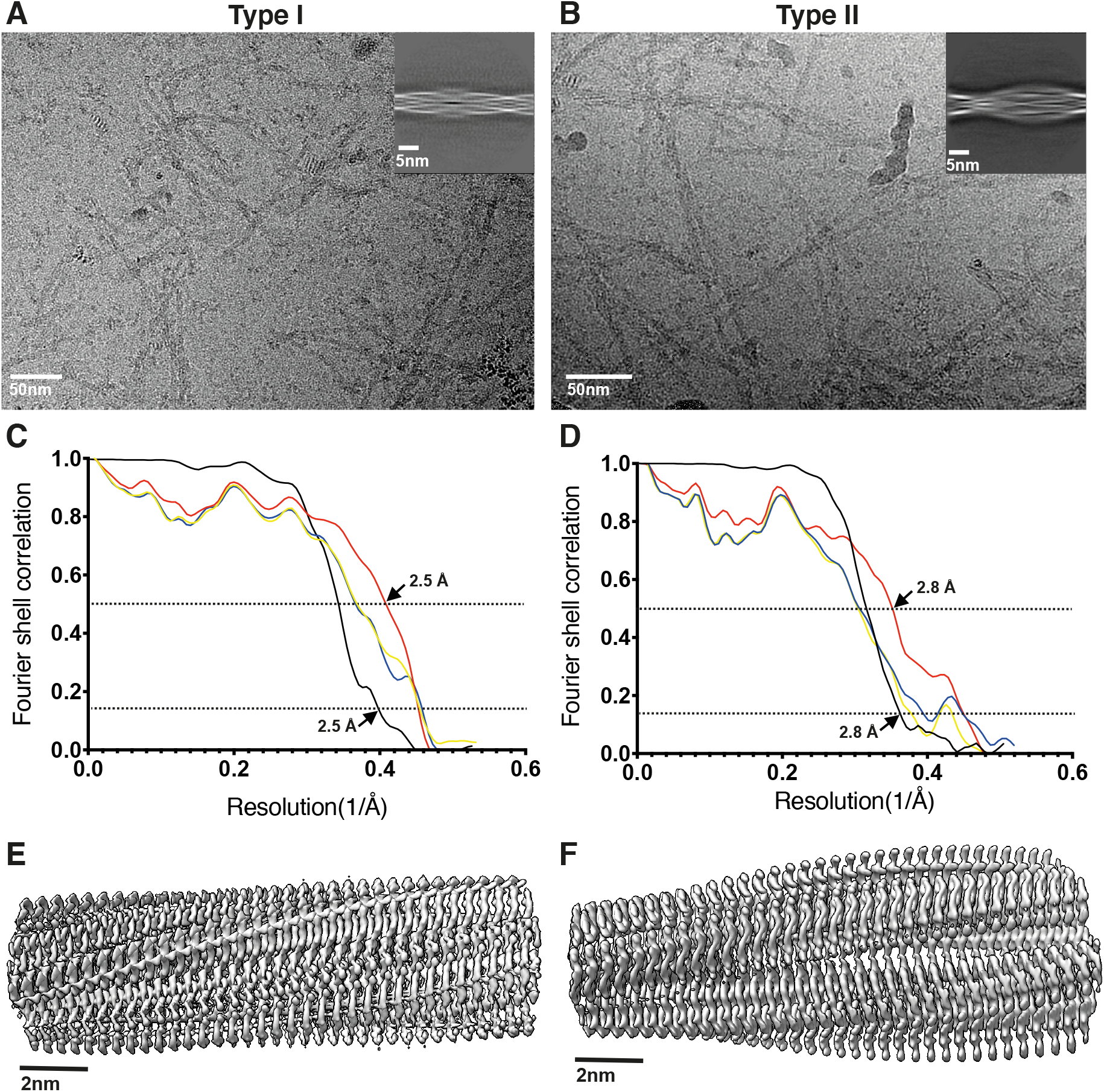
Cryo-EM micrographs and processing details. (**A,B**) Cryo-EM micrographs and 2D class averages of Type I and Type II Aβ42 filaments from Alzheimer’s disease case 1 (A) and the case of pathological aging (B). (**C,D**) Fourier shell correlation (FSC) curves for cryo-EM maps and structures of Type I and Type II Aβ42 filaments from Alzheimer’s disease case 1 (C) and the case of pathological aging (D). FSC curves for two independently refined cryo-EM half maps are shown in black; for the final refined atomic model against the final cryo-EM map in red; for the atomic model refined in the first half map against that half map in blue; and for the refined atomic model in the first half map against the other half map in yellow. (**E,F**) 3D reconstruction of Type I and Type II Aβ42 filaments from Alzheimer’s disease case 1 (E) and the case of pathological aging (F).

**Fig. S2.**
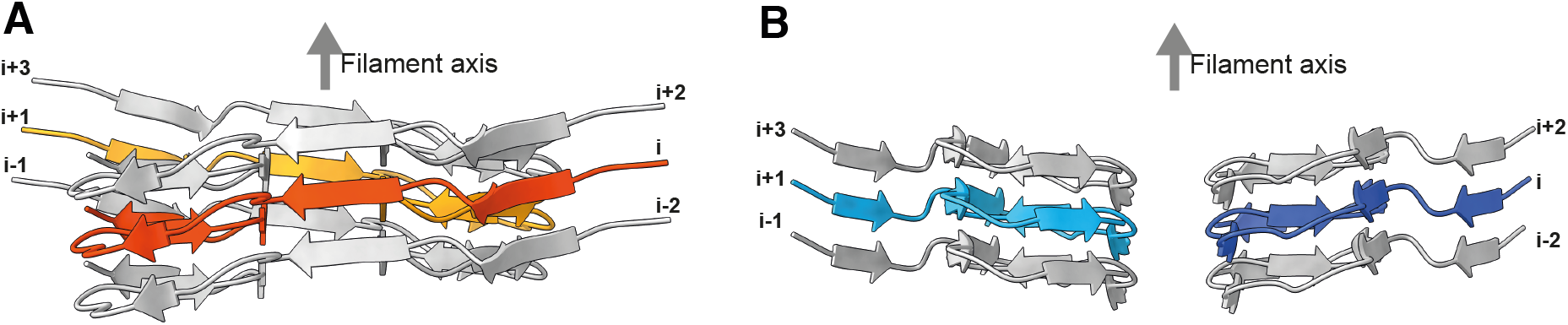
Protofilament symmetry packing of Type I and Type II Aβ42 filaments. (**A,B**) Rendered view of secondary structure elements in three successive rungs. The centre layer Type I monomer is shown in orange and yellow; the centre layer Type II monomer is shown in light and dark blue.

**Fig. S3.**
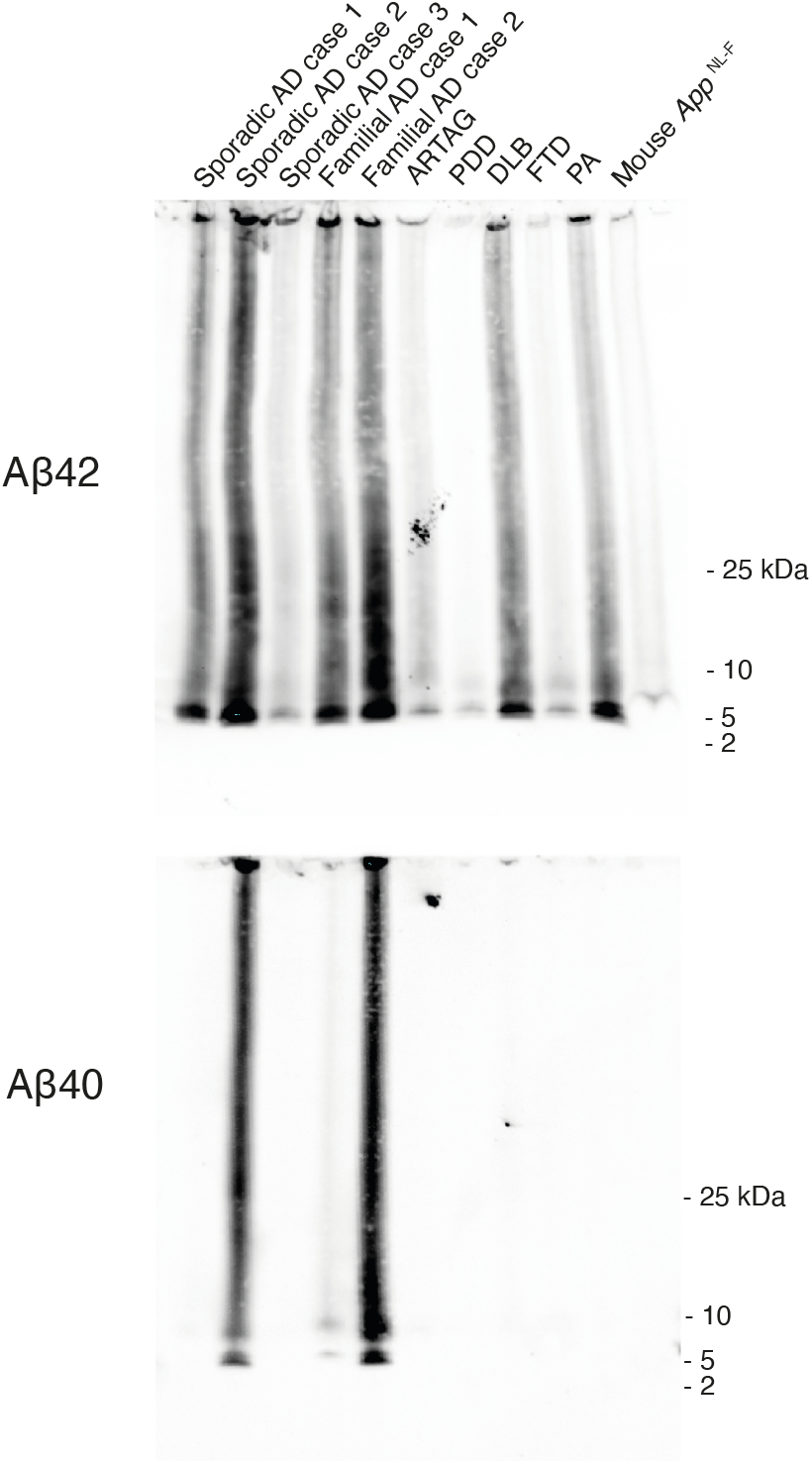
Immunoblot analysis. Labelling of Aβ42 and Aβ40 from the sarkosyl-insoluble fractions used for cryo-EM of five cases of Alzheimer’s disease (AD), a case of age-related tau astrogliopathy (ARTAG), a case of Parkinson’s disease dementia (PDD), a case of dementia with Lewy bodies (DLB), a case of frontotemporal dementia (FTD), a case of pathological aging (PA) and mouse line *App*^NL-F^. Monoclonal antibodies specific for Aβ42 and Aβ40 were used. For ARTAG, we used temporal cortex for immunoblotting and entorhinal cortex/hippocampus for cryo-EM. Aβ40, Aβ42 and their dimers are seen as individual bands. The smears and the immunoreactive material unable to enter the gel reflect the insolubility of Aβ assemblies.

**Fig. S4.**
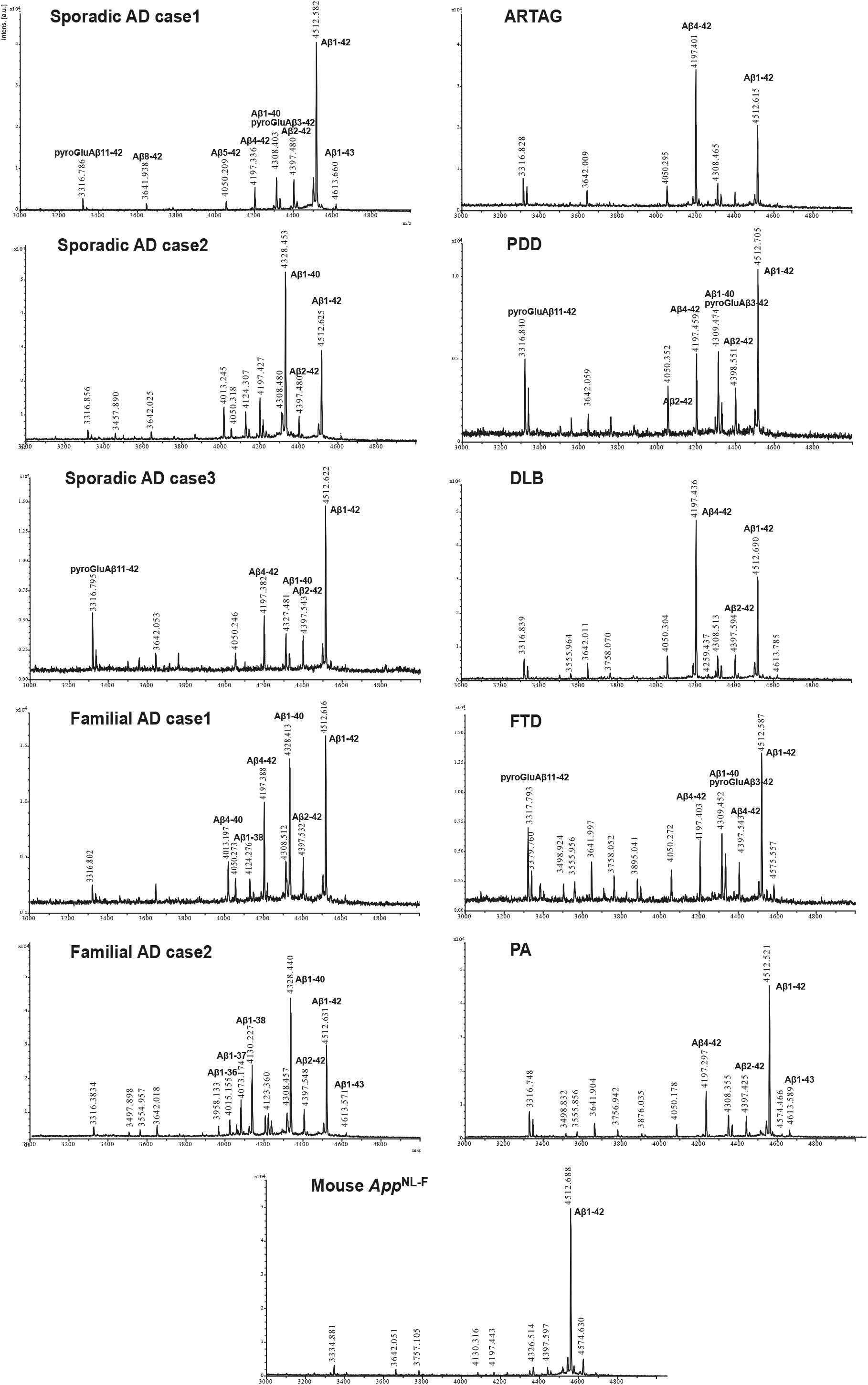
Mass spectrometric analysis. MALDI mass spectra for Aβ from sarkosyl-insoluble fractions used for cryo-EM of five cases of Alzheimer’s disease (AD), a case of age-related tau astrogliopathy (ARTAG), a case of Parkinson’s disease dementia (PDD), a case of dementia with Lewy bodies (DLB), a case of frontotemporal dementia (FTD), a case of pathological aging (PA) and mouse line *App*^NL-F^. For ARTAG, we used temporal cortex for mass spectrometry and entorhinal cortex/hippocampus for cryo-EM.

**Fig. S5.**
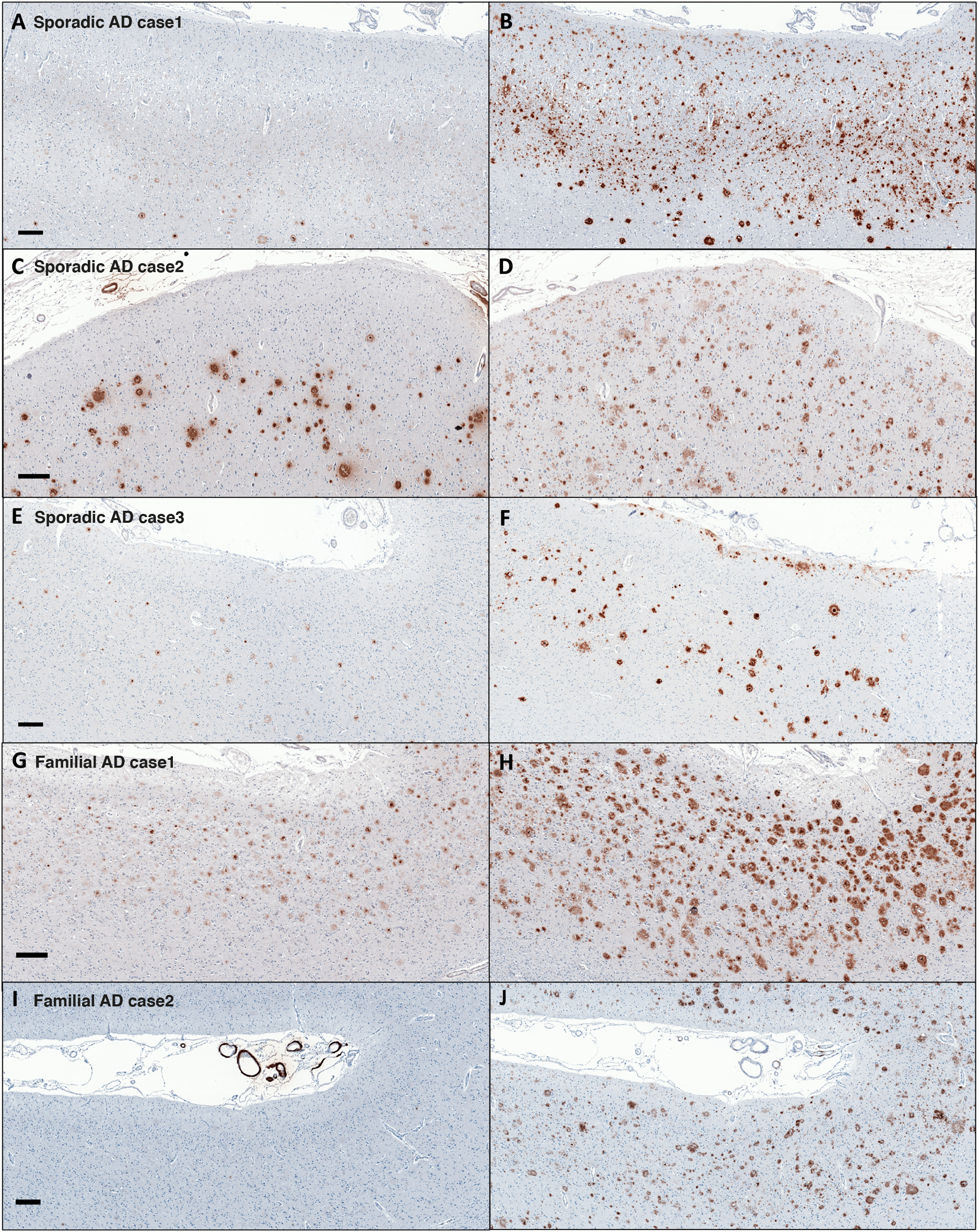
Sporadic cases 1-3 and familial cases 1-2 of Alzheimer’s disease: Immunohistochemistry for Aβ40 and Aβ42. Representative staining of the brain regions contralateral to those used for cryo-EM structure determination (see Methods), using monoclonal antibodies specific for Aβ40 (A,C,E,G,I) and Aβ42 (B,D,F,H,J). (**A,B**) Case 1 of sporadic Alzheimer’s disease; (**C,D**) Case 2 of sporadic Alzheimer’s disease; (**E,F**) Case 3 of sporadic Alzheimer’s disease; (**G,H**) Case 1 of familial Alzheimer’s disease; (**I,J**) Case 2 of familial Alzheimer’s disease. Scale bars, 200 μm.

**Fig. S6.**
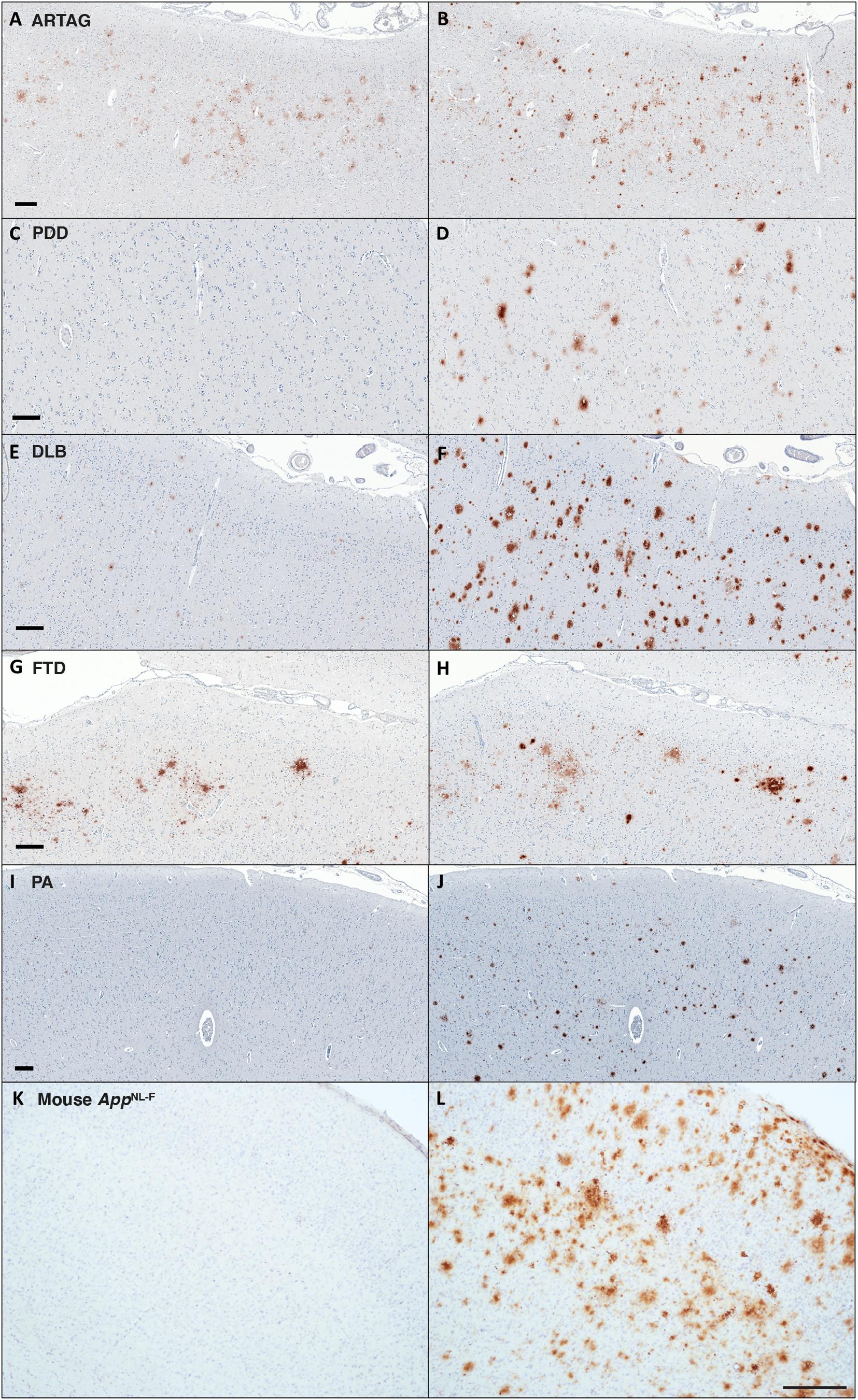
Human cases of ARTAG, PDD, DLB, FTD and PA, as well as frontal cortex from a brain of mouse line *App*^NL-F^: Immunohistochemistry for Aβ40 and Aβ42. Representative staining of the brain regions contralateral to those used for cryo-EM structure determination (see Methods), using monoclonal antibodies specific for Aβ40 (A,C,E,G,I,K) and Aβ42 (B,D,F,H,J,L). (A,B) ARTAG (Aging-related tau astrogliopathy); (C,D) PDD (Parkinson’s disease dementia); (E,F) DLB (Dementia with Lewy bodies); (G,H) FTD (Frontotemporal dementia); (I,J) PA (Pathological aging); (K,L) *App*^NL-F^ mouse (18-month-old). Scale bars, 200 μm.

**Fig. S7.**
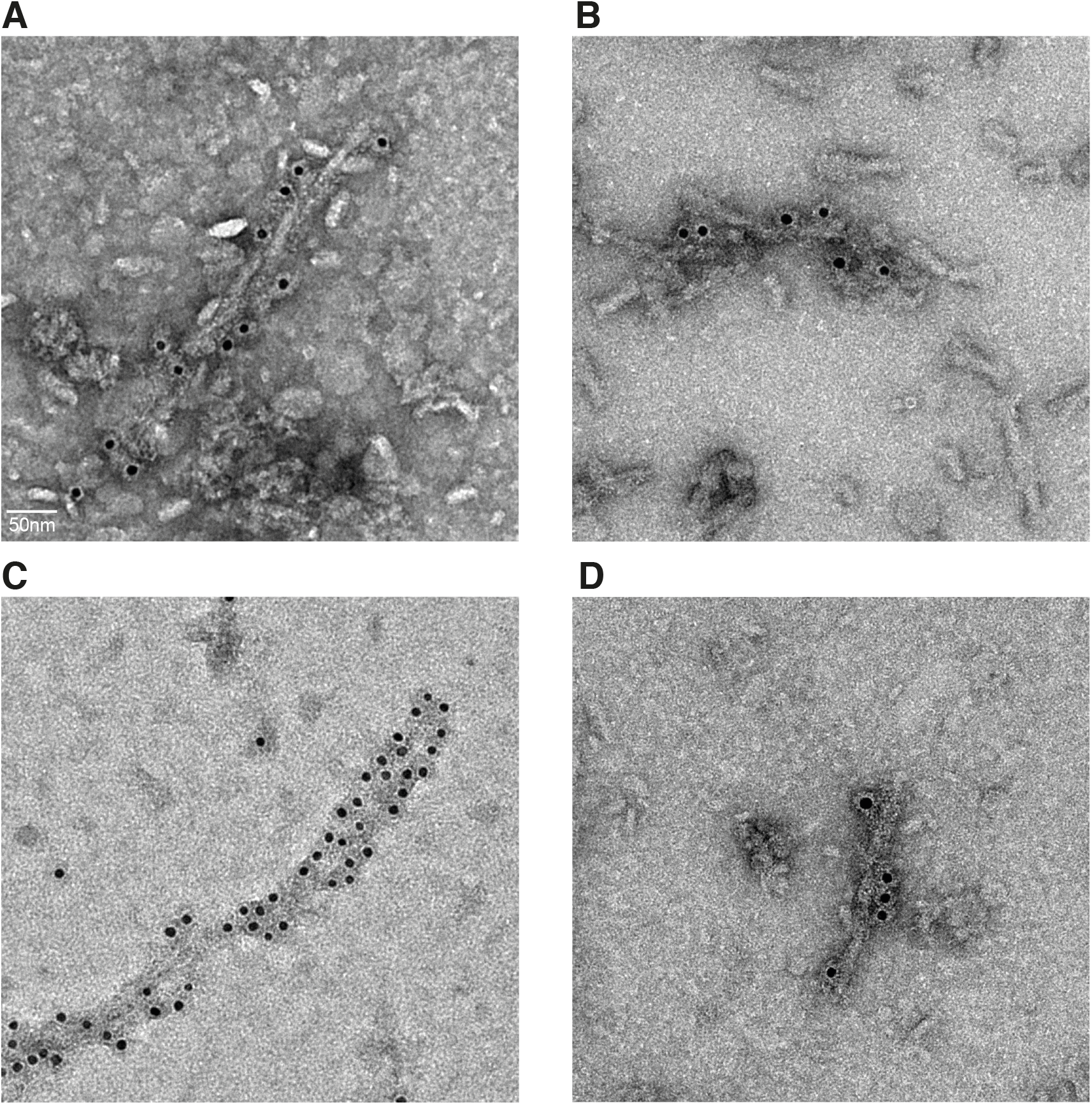
Immunogold negative-stain electron microscopy of Aβ filaments. (**A**) Sporadic Alzheimer’s disease case 1 with antibody D54D2. (**B**) Sporadic Alzheimer’s disease case 1 with antibody 1E11. (**C**) Case of pathological aging with antibody D54D2. (**D**) Case of pathological aging with antibody 1E11.

## References and Notes

1. J.M. Long, D.M. Holtzman, Alzheimer’s disease: An update on pathobiology and treatment strategies. Cell 179, 312–339 (2019).

2. J.A. Hardy, G.A. Higgins, Alzheimer’s disease: The amyloid cascade hypothesis. Science 256, 184–185 (1992).

3. C. Haass, C. Kaether, G. Thinakran, S. Sisodia, Trafficking and proteolytic processing of *APP*. Cold Spring Harb. Perspect. Med. 2, a006270 (2012).

4. N. Suzuki, T.T. Cheung, X.D. Cai, A. Odaka, L. Otvos, C. Eckman, T.E. Golde, S.G. Younkin, An increased percentage of long amyloid beta secreted by familial amyloid beta protein precursor (beta *APP*717) mutants. Science 264, 1336–1340 (1994).

5. D. Scheuner, C. Eckman, M. Jensen, X. Song, M. Citron, N. Suzuki, T.D. Bird, J. Hardy, M. Hutton, W. Kukull, E. Larson, E. Levy-Lahad, M. Viitanen, E. Peskind, P. Poorkaj, G. Schellenberg, R. Tanzi, W. Wasco, L. Lannfelt, D.J. Selkoe, S. Younkin, Secreted amyloid beta-protein similar to that in the senile plaques of Alzheimer’s disease is increased in vivo by the presenilin 1 and 2 and *APP* mutations linked to familial Alzheimer’s disease. Nature Med. 2, 864–870 (1996).

6. M. Citron, C. Vigo-Pelfrey, D.B. Teplow, C. Miller, D. Schenk, J. Johnston, B. Winblad, N. Venizelos, L. Lannfelt, D.J. Selkoe, Excessive production of amyloid-beta protein by peripheral cells of symptomatic and presymptomatic patients carrying the Swedish familial Alzheimer disease mutation. Proc. Natl. Acad. Sci. USA 91, 11993–11997 (1994).

7. M. Pagnon de la Vega, V. Giedraitis, W. Michno, L. Kilander, G. Güner, M. Zielinski, M. Löwenmark, R. Brundin, T. Danfors, L. Söderberg, I. Alazuloff, L.N.G. Nilsson, A. Erlandsson, D. Willbold, S.A. Müller, G.F. Schröder, J. Hanrieder, S.F. Lichtenthaler, L. Lannfelt, D. Sehlin, M. Ingelsson. The *Uppsala APP* deletion causes early onset autosomal dominant Alzheimer’s disease by altering *APP* processing and increasing amyloid β fibril formation. Sci. Transl. Med. 13, eabc6184 (2021).

8. T. Iwatsubo, A. Odaka, N. Suzuki, H. Mizusawa, N. Nukina, Y. Ihara, Visualization of Abeta42(43) and Abeta40 in senile plaques with end-specific Abeta monoclonals: Evidence that an initially deposited species is Abeta42(43). Neuron 13, 45–53 (1994).

9. A. Güntert, H. Döbeli, B. Bohrmann, High sensitivity analysis of amyloid-beta peptide composition in amyloid deposits from human and PS2APP mouse brain. Neuroscience 143, 461–475 (2006).

10. D.R. Thal, J. Walter, T.C. Saido, M. Fändrich, Neuropathology and biochemistry of Aβ and its aggregates in Alzheimer’s disease. Acta Neuropathol. 129, 167–182 (2015).

11. J. Attems, K.A. Jellinger, F. Lintner, Alzheimer’s disease pathology influences severity and topographical distribution of cerebral amyloid angiopathy. Acta Neuropathol. 110, 222–231 (2005).

12. M. Kollmer, W. Close, L. Funk, J. Rasmussen, A. Bsoul, A. Schierhorn, M. Schmidt, C.J. Sigurdson, M. Jucker, M. Fändrich, Cryo-EM structure and polymorphism of Aβ amyloid fibrils purified from Alzheimer’s brain tissue. Nature Commun. 10, 4760 (2019).

13. J.T. Jarrett, E.P. Berger, P.T. Lansbury, The carboxy terminus of the beta amyloid protein is critical for the seeding of amyloid formation: Implications for the pathogenesis of Alzheimer’s disease. Biochemistry 32, 4693–4697 (1993).

14. D.R. Thal, U. Rüb, M. Orantes, H. Braak, Phases of Abeta deposition in the human brain and its relevance for the development of Alzheimer’s disease. Neurology 58, 1791–1800 (2002).

15. M. Meyer-Luehmann, J. Coomaraswamy, T. Bolmont, S. Kaeser, C. Schaefer, E. Kilger, A. Neuenschwander, D. Abramowski, P. Frey, A.L. Jaton, J.M. Vigouret, P. Paganetti, D.M. Walsh, P.M. Matthews, J. Ghiso, M. Staufenbiel, L.C. Walker, M. Jucker, Exogenous induction of cerebral beta-amyloidogenesis is governed by agent and host. Science 313, 1781–1784 (2006).

16. H.H.C. Lau, M. Ingelsson, J.C. Watts, The existence of Aβ strains and their potential for driving phenotypic heterogeneity in Alzheimer’s disease. Acta Neuropathol. 142, 17–39 (2021).

17. Z. Jaunmuktane, S. Mead, M. Ellis, J.D.F. Wadsworth, A.J. Nicoll, J. Kenny, F. Launchbury, J. Linehan, A. Richard-Loendt, A.S. Walker, P. Rudge, J. Collinge, S. Brandner, Evidence for human transmission of amyloid-β pathology and cerebral amyloid angiopathy. Nature 525, 247–250 (2015).

18. K. Frontzek, M.I. Lutz, A. Aguzzi, G.G. Kovacs, H. Budka, Amyloid-β pathology and cerebral amyloid angiopathy are frequent in iatrogenic Creutzfeldt-Jakob disease after dural grafting. Swiss Med. Weekly 146, w14287 (2016).

19. Z. Jaunmuktane, A. Quaegebeur, R. Taipa, M. Viana-Baptista, R. Barbosa, C. Koriath, R. Sciot, S. Mead, S. Brandner, Evidence of amyloid-β cerebral angiopathy transmission through neurosurgery. Acta Neuropathol. 135, 671–679 (2018).

20. M.R. Sawaya, M.P. Hughes, J.A. Rodriguez, R. Riek, D.S. Eisenberg, The expanding amyloid family: Structure, stability, function, and pathogenesis. Cell 184, 4857–4873 (2021).

21. U. Ghosh, K.R. Thurber, W.-Y. Yau, R. Tycko, Molecular structure of a prevalent amyloid-β fibril polymorph from Alzheimer’s disease brain tissue. Proc. Natl. Acad. Sci. USA 118, e2023089118 (2021).

22. L. Gremer, D. Schölzel, C. Schenk, E. Reinartz, J. Labahn, R.B.G. Ravelli, M. Tusche, C. Lopez-Iglesias, W. Hoyer, H. Heise, D. Willbold, G.F. Schröder, Fibril structure of amyloid-β(1-42) by cryo-electron microscopy. Science 358, 116–119 (2017).

23. Y. Xiao, B. Ma, D. McElheny, S. Parthasarathy, F. Long, M. Hoshi, R. Nussinov, Y. Ishii, Aβ(1-42) fibril structure illuminates self-recognition and replication of amyloid in Alzheimer’s disease. Nature Struct. Mol. Biol. 22, 499–505 (2015).

24. M.T. Colvin, R.l. Silvers, Q.Z. Ni, T.V. Can, I. Sergeyev, M. Rosay, K.J. Donovan, B. Michael, J. Wall, S. Linse, R.C. Griffin, Atomic resolution structure of monomorphic Aβ42 amyloid fibrils. J. Am. Chem. Soc. 138, 9663–9674 (2016).

25. M.A. Wälti, F. Ravotti, H. Arai, C.G. Glabe, J.S. Wall, A. Böckmann, P. Güntert, B.H. Meier, R. Riek, Atomic resolution structure of a disease-relevant Aβ(1-42) amyloid fibril. Proc. Natl. Acad. Sci. USA 113, E4976–E4984 (2016).

26. A.K. Schütz, T. Vagt, M. Huber, O.Y. Ovchinnikova, R. Cadalbert, J. Wall, P. Güntert, A. Böckmann, R. Glockshuber, B.H. Meier, Atomic-resolution three-dimensional structure of amyloid β fibrils bearing the Osaka mutation. Angew. Chem. Int. Ed. 54, 331–335 (2015).

27. R. Vassar, P.H. Kuhn, C. Haass, M.E. Kennedy, L. Rajendran, P.C. Wong, S.F. Lichtenthaler, Function, therapeutic potential and cell biology of BACE proteases: Current status and future prospects. J. Neurochem. 130, 4–28 (2014).

28. B. Falcon, W. Zhang, M. Schweighauser, A.G. Murzin, R. Vidal, H.J. Garringer, B. Ghetti, S.H.W. Scheres, M. Goedert, Tau filaments from multiple cases of sporadic and inherited Alzheimer’s disease adopt a common fold. Acta Neuropathol. 146, 699–708 (2018).

29. A.W.P. Fitzpatrick, B. Falcon, S. He, A.G. Murzin, G. Murshudov, H.J. Garringer, R.A. Crowther, B. Ghetti, M. Goedert, S.H.W. Scheres, Cryo-EM structures of tau filaments from Alzheimer’s disease. Nature 547, 185–190 (2017).

30. W. Zhang, B. Falcon, A.G. Murzin, J. Fan, R.A. Crowther, M. Goedert, S.H.W. Scheres, Heparin-induced tau filaments are polymorphic and differ from those of Alzheimer’s and Pick’s diseases. eLife 8, e43584 (2019).

31. M. Schweighauser, Y. Shi, A. Tarutani, F. Kametani, A.G. Murzin, B. Ghetti, T. Matsubara, T. Tomita, T. Ando, K. Hasegawa, S. Murayama, M. Yoshida, M. Hasegawa, S.H.W. Scheres, M. Goedert, Structures of α-synuclein filaments from multiple system atrophy. Nature 585, 464–469 (2020).

32. S. Lövestam, M. Schweighauser, T. Matsubara, S. Murayama, T. Tomita, T. Ando, K. Hasegawa, M. Yoshida, A. Tarutani, M. Hasegawa, M. Goedert, S.H.W. Scheres, Seeded assembly in vitro does not replicate the structures of α-synuclein filaments from multiple system atrophy. FEBS Open Bio 11, 999–1013 (2021).

33. T. Saito, Y. Matsuba, N. Mihira, J. Takano, P. Nilsson, S. Itohara, N. Iwata, T.C. Saido, Single *App* knock-in mouse models of Alzheimer’s disease. Nature Neurosci. 17, 661–663 (2014).

34. Y. Shi, W. Zhang, Y. Yang, A.G. Murzin, B. Falcon, A. Kotecha, M. van Beers, A. Tarutani, F. Kametani, H.J. Garringer, R. Vidal, G.I. Hallinan, T. Lashley, Y. Saito, S. Murayama, M. Yoshida, H. Tanaka, A. Kakita, T. Ikeuchi, A.C. Robinson, D.M.A. Mann, G.G. Kovacs, T. Revesz, B. Ghetti, M. Hasegawa, M. Goedert, S.H.W. Scheres, Structure-based classification of tauopathies. Nature, in press. Doi: 10.1038/s41586-021-03911-7.

35. C.D. Chen, N. Joseph-Mathurin, N. Sinha, A. Zhou, Y. Li, K. Friedrichsen, A. McCullough, E.E. Franklin, R. Hornbeck, B. Gordon, V. Sharma, C. Cruchaga, A. Goate, C. Karch, E. McDade, C. Xiong, R.J. Bateman, B. Ghetti, J.M. Ringman, J. Chhatwal, C.L. Masters, C. McLean, T. Lashley, Y. Su, R. Koeppe, C. Jack, W.E. Klunk, J.C. Morris, R.J. Perrin, N.J. Cairns, T.L.S. Benzinger, Comparing amyloid-β plaque burden with antemortem PiB PET in autosomal dominant and late-onset Alzheimer’s disease. Acta Neuropathol. 142, 689–708 (2021).

36. S.F. Lichtenthaler, D. Beher, H.S. Grimm, R. Wang, M.S. Shearman, C.L. Masters, K. Beyreuther, The intramembrane cleavage site of the amyloid precursor protein depends on the length of its transmembrane domain. Proc. Natl. Acad. Sci. USA 99, 1365–1370 (2002).

37. C. Guardia-Laguarta, M. Pera, J. Clarimón, J.L. Molinuevo, R. Sánchez-Valle, A. Lladó, M. Coma, T. Gómez-Isla, R. Blesa, I. Ferrer, A. Lleó, Clinical, neuropathologic, and biochemical profile of the amyloid precursor protein I716F mutation. J. Neuropathol. Exp. Neurol. 69, 553–59 (2010). Doi: 10.1097/NEN.0b013e3181c6b84.

38. J. Rasmussen, J. Mahler, N. Beschorner, S.A. Kaeser, L.M. Häsler, F. Baumann, S. Nyström, E. Portelius, K. Blennow, T. Lashley, N.C. Fox, D. Sepulveda-Falla, M. Glatzel, A.L. Oblak, B. Ghetti, K.P.R. Nilsson, P. Hammerström, M. Staufenbiel, L.C. Walker, M. Jucker, Amyloid polymorphisms constitute distinct clouds of conformational variants in different etiological subtypes of Alzheimer’s disease. Proc. Natl. Acad. Sci. USA 114, 13018–13023 (2017).

39. J.X. Lu, W. Qiang, W.M. Yau, C.D. Schwieters, S.C. Meredith, R. Tycko, Molecular structure of β-amyloid fibrils in Alzheimer’s disease brain tissue. Cell 154, 1257–1268 (2013).

40. M.E. Murray, D.W. Dickson, Is pathological aging a successful resistance against amyloid-beta or preclinical Alzheimer’s disease? Alzheimer’s Res. Ther. 6, 24 (2014).

41. J.R. Murrell, M. Farlow, B. Ghetti, M.D. Benson, A mutation in the amyloid precursor protein associated with hereditary Alzheimer’s disease. Science 254, 97–99 (1991).

42. C.L. Maarouf, I.D. Daugs, S. Spina, R. Vidal, T.A. Kokjohn, R.L. Patton, W.M. Kalback, D.C. Luehrs, D.G. Walker, E.M. Castaño, T.G. Beach, B. Ghetti, A.E. Roher, Histopathological and molecular heterogeneity among individuals with dementia associated with presenilin mutations. Mol. Neurodeg. 3, 20 (2008).

43. S. Klotz, P. Fischer, M. Hinterberger, G. Ricken, S., Hönigschnabl, E. Gelpi, G.G. Kovacs, Multiple system aging-related tau astrogliopathy with complex proteinopathy in an oligosymptomatic octogenarian. Neuropathology 41, 72–83 (2021).

44. S. Spina, M.R. Farlow, F.W. Unverzagt, D.A. Kareken, J.R. Murrell, G. Fraser, F. Epperson, R.A. Crowther, M.G. Spillantini, M. Goedert, B. Ghetti, The tauopathy associated with mutation +3 in intron 10 of Tau: Characterization of the MSTD family. Brain 131, 72–89 (2008).

45. Näslund J. Schierhorn A, Hellman U, Lannfelt L, Roses AD, Tjernberg LO, Silberring J, Gandy SE, Winblad B, Greengard P, Norstedt C, Terenius L, Relative abundance of Alzheimer Aβ amyloid peptide variants in Alzheimer disease and normal aging. Proc. Natl. Acad. Sci. USA 91, 8378–8382 (1994).

46. Kakuda N, Miyasaka T, Iwasaki N, Nirasawa T, Wada-Kakuda S, Takahashi-Fujigasaki J, Murayama S, Ihara Y, Ikegawa M, Distinct deposition of amyloid-β species in brains with Alzheimer’s disease pathology visualized with MALDI imaging mass spectrometry. Acta Neuropathol. Commun. 5, 73 (2017).

47. J.L. Farlow, L.A. Robak, K. Hetrick, K. Bowling, E. Boerwinkle, Z.H. Coban-Akdemir, T. Gambin, R.A. Gibbs, S. Gu, P. Jain, J. Jankovic, S. Jhangiani, K. Kaw, D. Lai, H. Lin, H. Ling, Y. Liu, J.R. Lupski, D. Muzny, P. Porter, E. Pugh, J. White, K. Doheny, R.M. Myers, J.M. Shulman, T. Foroud, Whole-exome sequencing in familial Parkinson disease. JAMA Neurol. 73, 68–75 (2016).

48. M. Goedert, M.G. Spillantini, N.J. Cairns, R.A. Crowther, Tau proteins of Alzheimer paired helical filaments: Abnormal phosphorylation of all six brain isoforms. Neuron 8, 159–168 (1992).

49. S.Q. Zheng, E. Palovcak, J.P. Armache, K.A. Verba, Y. Cheng, D.A. Agard, MotionCor2: Anisotropic correction of beam-induced motion for improved cryoelectron microscopy. Nature Meth. 14, 331–332 (2017).

50. K. Zhang, Gctf: Real-time CTF determination and correction. J. Struct. Biol. 193, 1–12 (2016).

51. S. He, S.H.W. Scheres, Helical reconstruction in RELION. J. Struct. Biol. 198, 163–176 (2017).

52. J. Zivanov, T. Nakane, B.O. Forsberg, D. Kimanius, W.J. Hagen, E. Lindahl, S.H.W. Scheres, New tools for automated high-resolution cryo-EM structure determination in RELION-3. eLife 7, e42166 (2018).

53. S.H.W. Scheres, Amyloid structure determination in RELION-3.1. Acta Crystallogr. D Struct. Biol. 76, 94–101 (2020).

54. J. Zivanov, T. Nakane, S.H.W. Scheres, A Bayesian approach to beam-induced motion correction in cryo-EM single-particle analysis. IUCrJ 6, 5–17 (2019).

55. J. Zivanov, T. Nakane, S.H.W. Scheres, Estimation of high-order aberrations and anisotropic magnification from cryo-EM data sets in RELION-3.1. IUCrJ 7, 253–267 (2020).

56. S. Chen, G. McMullan, A.R. Faruqi, G.N. Murshudov, J.M. Short, S.H.W. Scheres, R. Henderson, High-resolution noise substitution to measure overfitting and validate resolution in 3D structure determination by single particle electron cryomicroscopy. Ultramicroscopy 135, 24–35 (2013).

57. R.A. Warmack, D.R. Boyer, C.-T. Zee, L.S. Richards, M.R. Sawaya, D. Cascio, T. Gonen, D.S. Eisenberg, S.G. Clarke, Structure of amyloid-β (20-34) with Alzheimer’s-associated isomerization at Asp23 reveals a distinct protofilament interface. Nature Commun. 10: 2257 (2019).

58. A. Casañal, B. Lohkamp, P. Emsley, Current developments in Coot for macromolecular model building of electron cryo-microscopy and crystallographic data. Protein Sci. 29, 1069–1078 (2020).

59. C.J. Williams, J.J. Headd, N.W. Moriarty, M.G. Prisant, L.L. Videau, L.N. Deis, V. Verma, D.A. Keedy, B.J. Hintze, V.B. Chen, S. Jain, S.M. Lewis, W.B. Arendall, J. Snoeyink, P.D. Adams, S.C. Lovell, J.S. Richardson, D.C. Richardson, MolProbity: More and better reference data for improved all-atom structure validation. Protein Sci. 27, 293–315 (2018).

60. A. Brown, F. Long, R.A. Nicholls, J. Toots, P. Emsley, G. Murshudov, Tools for macromolecular model building and refinement into electron cryo-microscopy reconstructions. Acta Crystallogr. D Biol. Crystallogr. 71, 136–153 (2015).

61. P.V. Afonine, B.K. Poon, R.J. Read, O.V. Sobolev, T.C. Terwilliger, A. Urzhumtsev, P.D. Adams, Real-space refinement in PHENIX for cryo-EM and crystallography. Acta Crystallogr. D Struct. Biol. 74, 531–544 (2018).

62. E.F. Pettersen, T.D. Goddard, C.C. Huang, E.C. Meng, G.S. Couch, T.I. Croll, J.H. Morris, T.E. Ferrin, UCSF ChimeraX: Structure visualization for researchers, educators, and developers. Protein Sci. 30, 70–82 (2021).

